# Joint nonparametric coalescent inference of mutation spectrum history and demography

**DOI:** 10.1101/2020.06.16.153452

**Authors:** William S. DeWitt, Kameron Decker Harris, Kelley Harris

## Abstract

Booming and busting populations modulate the accumulation of genetic diversity, encoding histories of living populations in present-day variation. Many methods exist to decode these histories, and all must make strong model assumptions. It is typical to assume that mutations accumulate uniformly across the genome at a constant rate that does not vary between closely related populations. However, recent work shows that mutational processes in human and great ape populations vary across genomic regions and evolve over time. This perturbs the *mutation spectrum*: the relative mutation rates in different local nucleotide contexts. Here, we develop theoretical tools in the framework of Kingman’s coalescent to accommodate mutation spectrum dynamics. We describe mushi: a method to perform fast, nonparametric joint inference of demographic and mutation spectrum histories from allele frequency data. We use mushi to reconstruct trajectories of effective population size and mutation spectrum divergence between human populations, identify mutation signatures and their dynamics in different human populations, and produce more accurate time calibration for a previously-reported mutational pulse in the ancestors of Europeans. We show that mutation spectrum histories can be productively incorporated in a well-studied theoretical setting, and rigorously inferred from genomic variation data like other features of evolutionary history.

## Introduction

Over the past decade, population geneticists have developed many sophisticated methods for inferring population demography, and have consistently found that simple, isolated populations of constant size are far from the norm. Population expansions and founder events, as well as migration between species and geographic regions, have been inferred from virtually all high resolution genetic datasets that have been generated, and we now recognize that inferring these non-equilibrium demographies is often essential for understanding histories of adaptation and global migration. Population genetics has uncovered many features of human history that were once virtually unknowable by other means [1], revealing a complex series of migrations, population replacements, and admixture networks among human groups and extinct hominoids. Related analyses of genetic variation have also shown that ancestral human populations differed from one another at the biochemical level, inheriting systematically different patterns of DNA damage. It is not known how many of these differences were genetically encoded as opposed to environmentally induced, but either type of perturbation has the potential to complicate the task of inferring population history from genetic variation.

The process of germline mutation is the writing mechanism that records signatures of demographic events in genomes, so its influence on modern genomic variation is similar in importance to the demographic histories themselves. Demographic inference methods can model complex population splits, migration, and admixture, and while some have the potential to accommodate various functional forms for *N* (*t*), mutation has long received a comparatively simple treatment. Usually, a single mutation rate parameter *μ* is assumed to apply at all loci, in all individuals, and at all times. It may then be regarded as a nuisance parameter needed for time calibration of models where time is measured in dimensionless Wright-Fisher generations (i.e. units of 2*N*). *De novo* mutation rates in humans can be measured by parent-child trio sequencing studies, while for other species it is typical to use a phylogenetically calibrated mutation rate parameter, and the accuracy of these often uncertain estimates places a fundamental limit on the precision of inferred parameters such as times of admixture and population divergence.

Although modern methods for inferring demography from genetic data tend to assume a mutational process that is simple and unchanging, mutation rate evolution has long been a subject of study in population genetics. Soon after Haldane developed equilibrium theory for alleles in mutation-selection balance and used this to provide the first principled estimate of the human mutation rate by studying hemophilia incidence [2, 3], Kimura began to consider how *mutator alleles*—i.e. genetic modifiers of the mutation rate—had the potential to optimize mutation rates by balancing adaptive response to environmental changes against increasing genetic load [4]. Kimura recognized the tendency of mutators to escape their deleterious consequences via recombination away from new mutations that they help create, and therefore deduced that rising mutation rates might be a deleterious consequence of increasing reliance on sexual reproduction. The *drift-barrier hypothesis* of Lynch et al. expands upon this idea by considering the effect of genetic drift on mutation rate optima. Population bottlenecks and low effective population size ultimately limit the ability of a population to evolve toward an optimum of high replication fidelity, as the efficiency of selection against mutator alleles increases with *N* [5].

Growing evidence indicates that germline mutation is a dynamic process that evolves over both interspecific and population time scales. The rate of this evolution has the potential to be highly pleiotropic; influenced by replication machinery polymorphisms as well as life history, mutagenic exposures, and genomic features such as repeats and epigenetic marks. Mutation rates among great apes appear to have declined along the lineage leading to humans—a phenomenon called the *hominoid slowdown* [6, 7]—, showing that mutation rate evolution between species distorts phylogenetic time calibration. At the level of single generations, children of older parents receive more germline mutations, especially from older fathers. Replicative errors in spermatogenesis add *≈* 1 additional expected mutation per year of paternal age, and the repair efficiency of spermatocyte DNA damage declines with age [8]. This *parental age effect* [9] means that sex-specific life history traits can influence mutagenesis at the population level. The first few embryonic cell divisions are more error prone than others [10], further demonstrating that not all cell divisions are clock-like. These phenomena show that the accumulation of mutations is complexly coupled to other biological processes.

A complex and polymorphic mutation process also reveals itself in associations with genomic position and local nucleotide context. The rate of C*→*T transitions is elevated at methylated CpG sites due to spontaneous deamination [11, 12]. GC-biased gene conversion (gbGC) refers to the tendency of stronger-binding GC alleles to overwrite AT alleles during homologous recombination [13, 14]. This biased non-Mendelian segregation pattern is tantamount to selection for weak-to-strong mutations from AT to GC, and can create new, sequence-biased mutations when non-allelic gene conversion transfers variation between paralogous genomic regions.

It is difficult to disentangle past changes in mutation rate from past changes in effective population size, which can change the rate of nucleotide substitution even when mutation rate stays constant. However, evolution of the mutation process can be indirectly detected by measuring its effects on the *mutation spectrum*: the relative mutation rates among different local nucleotide contexts. Hwang and Green [11] modeled the triplet context-dependence of the substitution process in a mammalian phylogeny, finding varying contributions from replication errors, cytosine deamination, and biased gene conversion. Many cancers have elevated mutation rates due to different failure points in the DNA repair process, and these differences cause hypermutation in different sets of triplet sequence motifs [15, 16]. Harris and Pritchard [17, 18] demonstrated the power of examining the same triplet-based spectrum in an evolutionary context, and counted single nucleotide variants in each triplet mutation type as a proxy for mutational input from each individual’s history. Human triplet spectra distinctly cluster according to continental ancestry group, and evidence of historical pulses in mutation activity (or suppression of repair) has been found in the distribution of allele frequencies in certain mutation types. Mathiesen et al. studied similar mutation signatures in rare human variants [19], and clarified alternative non-mutational hypotheses for their origin, including population differences in demography, patterns of selection, recombination, or recombination-associated processes such as gene conversion. Rare variants in large cohorts serve as a proxy for recent de novo mutations, and they reveal mutational signatures of replication timing, recombination, and sex differences in repair processes [20, 21].

Emerging from the literature is a picture of a mutation process evolving within and between populations, anchored to genomic features and accented by spectra of local nucleotide context. If probabilistic models of population genetic processes are to keep pace with these empirical findings, mutation deserves a richer treatment in state-of-the-art inference tools. In this paper, we build on classical theoretical tools to introduce fast nonparametric inference of population-level *mutation spectrum history* (MuSH)—the relative mutation rate in different local nucleotide contexts across time—alongside inference of demographic history. Whereas previous work has demonstrated mutation spectrum evolution using phenomenological statistics on modern variation, we shift perspective to focus on inference of the MuSH, which we model on the same footing as demography.

Demographic inference requires us to invert the map that takes population history to the patterns of genetic diversity observable today. This task is often simplified by first compressing these genetic diversity data into a summary statistic such as the *sample frequency spectrum* (SFS), the distribution of derived allele frequencies among sampled haplotypes. The SFS is a well-studied population genetic summary statistic that is sensitive to demographic history. Unfortunately, inverting the map from demographic history to SFS is a notoriously ill-posed problem, in that many different population histories can have identical expected SFS [22, 23, 24, 25, 26]. One way to deal with the ill-posedness of demographic inference (and other inverse problems) is to specify a parametric model. This is done by allowing a small number of constant or exponential epochs whose location and scale parameters are optimized to recapitulate the patterns observed in genomic data. An alternative is to allow a more general space of solutions, but to *regularize* by penalizing histories that contain features deemed biologically unrealistic (e.g. high frequency population size oscillations). Both approaches shrink the set of feasible solutions to the inverse problem so that it becomes well-posed, and can be thought of as leveraging prior knowledge. In particular, the penalization approach leverages knowledge about the granularity of generations in the discrete-time reproductive models that the continuous-time coalescent only approximates.

In this paper, we extend a coalescent framework for demographic inference to accommodate inference of the MuSH from a SFS that is resolved into different local *k*-mer nucleotide contexts. This is a richer summary statistic that we call the *k*-SFS, where e.g. *k* = 3 means triplet context. We show using coalescent theory that the *k*-SFS is related to the MuSH by a linear transformation, while depending non-linearly on the demographic history. We jointly infer both demographic history and MuSH using optimization, where the cost that we minimize balances a data fitting term, which employs the forward map from coalescent theory, along with a regularization term that favors smooth solutions with low complexity. Our open-source software mushi (mutation spectrum history inference) is available at https://harrispopgen.github.io/mushi as a Python package alongside computational notebooks that both demonstrate its use and reproduce the results of this paper. Using default settings and modest hardware, mushi takes only a few seconds to infer histories from population-scale sample frequency data.

The recovered MuSH is a rich object that illuminates both standard and previously hidden dimensions of population history. Various biological questions about evolution of the mutation process may be addressed by computing MuSH summary statistics, both intrapopulation (patterns within a single MuSH) and interpopulation (comparisons between MuSHs). After validating with data simulated under known histories, we use mushi to independently infer histories for each of the 26 populations (from 5 super-populations defined by continental ancestry) from the 1000 Genomes Project (1KG) [27]. We demonstrate that mushi is a powerful tool for demographic inference that has several advantages over existing demographic inference methods, then go on to describe the newly illuminated features of human mutation spectrum evolution.

We recover accurately timed demographic features that are robust to regularization parameter choices, including the out-of-Africa event (OOA) and the more recent bottleneck in the ancestors of modern Finns, and we find that effective population sizes converge ancestrally within each super-population, despite being inferred independently. Decomposing human MuSH into principal mutation signatures varying through time in each population, we find evidence of global divergence in the mutation process impacting many mutation types, and recapitulate trees of population and super-population relatedness. Finally, we revisit the timing of a previously reported ancient pulse of elevated TCC*→*TTC mutation rate, active primarily in the ancestors of Europeans, and absent in East Asians [17, 18, 28]. We find that the extent of the pulse into the ancient past is exquisitely sensitive to the choice of demographic history model, and that our best-fitting demographic model yields a pulse timing that is significantly older than previously thought, seemingly arising before the divergence time of East Asians and Europeans.

With mushi we can quickly reconstruct demographic history and MuSH without strong model specification requirements. This adds a new approach to the toolbox for researchers interested only in demographic inference. For researchers studying the mutation spectrum, accurate demographic history is essential if time calibration of events in mutation history are sought. Thus we expect that jointly modeling demography and mutation spectrum history will be an important tool for studying complex histories of mutational processes in population genetics.

## Model Summary

### Augmenting the SFS with nucleotide context information

The standard sample frequency spectrum (SFS) is a summary statistic of population variation that counts variants partitioned by the number of sampled individuals who carry the derived allele. Since rare variants tend to be younger than common variants, this summary preserves considerable information about the distribution of allele age, which can reflect temporal variation in either the mutation rate or the intensity of genetic drift. To disentangle these two causal factors, we leverage the fact that genetic drift affects all mutations uniformly, whereas the mutation rate is more likely to exhibit patterns of change that differ between genomic sequence contexts.

We could choose to partition mutations by any desired genomic characteristics, including their presence in epigenetically modified functional genomic regions, but in this work we focus on classifying mutations by their derived allele and the ancestral *k*-mer nucleotide contexts in which they occur, with *k* odd and the variant occupying the central position of the motif. There are *κ* = 2 × 3 × 4^*k−*1^ mutation types after collapsing by strand symmetry. For example, when *k* = 3 there are *κ* = 96 triplet mutation types, of which TCC*→*TTC is one. For a sample of *n* genomes, the standard SFS is an (*n −* 1)-dimensional vector of the number of variants present in exactly *i* genomes, with *i* ranging from 1 to *n −* 1. In contrast, the *k*-SFS is an (*n −* 1) *× κ*-dimensional matrix, where the (*i, j*)-th entry is the number of variants present in exactly *i* individuals that stem from mutations of type *j* (from one particular *k*-mer to another).

Our goal is to jointly infer demographic history and MuSH by efficiently searching for histories that optimize a composite likelihood of an observed *k*-SFS data matrix **X**. This requires computing **Ξ***≡* 𝔼[**X**], the expected *k*-SFS as a function of effective population size and context-dependent mutation intensity over time. Our main theoretical result, Theorem 1 in the Methods, shows that **Ξ** is a linear functional of the *κ*-element mutation spectrum history ****μ****(*t*) given the haploid effective population size history *η*(*t*) (where *η*(*t*) = 2*N* (*t*) for diploid populations): 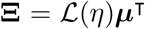 Figure 1a sketches the generation of a sampled *k*-SFS matrix **X**in a toy setting of *n* = 4 sampled haplotypes, 3 mutation types, and a fixed genealogy. Figure 1b clarifies the action of the linear operator 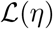.

**Figure 1:**
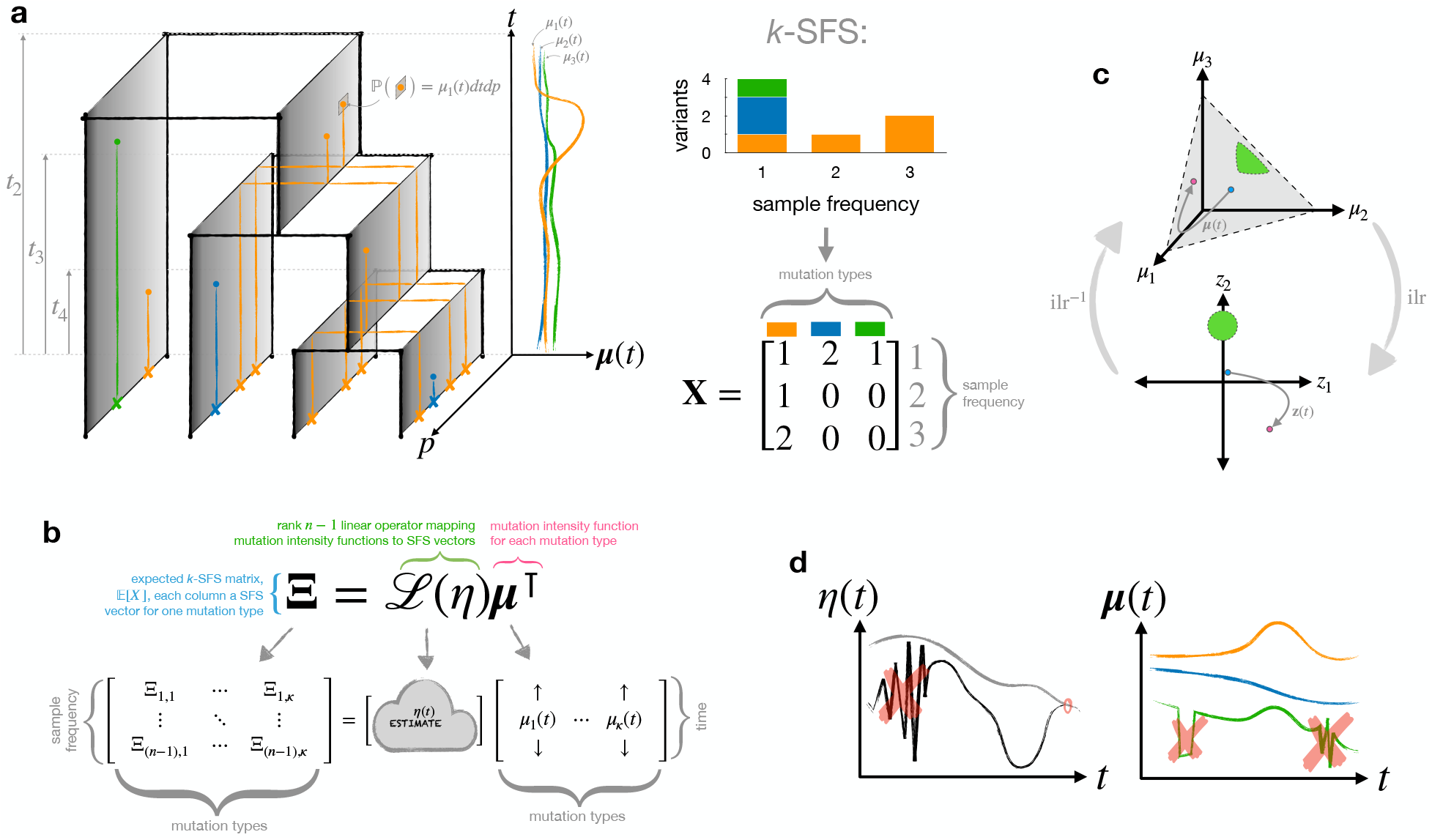
Mutation spectrum history and demography are encoded in the *k*-SFS as joint inverse problems. **a.** A schematic of a marked Poisson process with *n* = 4 sampled haplotypes is conditioned on coalescent times *t*_4_*, t*_3_*, t*_2_. The mutation spectrum history ***μ***(*t*) = [*μ*_1_(*t*) *μ*_2_(*t*) *μ*_3_(*t*)]^T^ shows just three mutation types (colors). Dots indicate mutation events placed by time *t*, genomic position *p*, and coalescent line (which are depicted as extruded in the genomic coordinate axis, grey sheets). The probability that a mutation of type *i* occurs in a differential time interval *dt* and genomic interval *dp* on a given line is proportional to the instantaneous mutation intensity *μ_i_*(*t*). The crosses on the sampled haplotypes indicate segregating variants of each mutation type. The sampled *k*-SFS data is shown as a stacked histogram (top right), and in matrix form (bottom right). **b.** Unpacking the forward map from MuSH ***μ***(*t*) and demography *η*(*t*) to expected *k*-SFS **Ξ**. **c.** Schematic of the isometric log ratio transform for compositional data, which maps the simplex (top) to a Euclidean space (bottom) in which optimization is more easily performed. **d.** Schematic of regularization concepts for inferring *η*(*t*) and ***μ***(*t*). Complex oscillations in time are penalized, as is the number of independent mutation spectrum components, and ancestral convergence may be encouraged.

### Using regularization to select parsimonious population histories

Even ordinary demographic inference—the recovery of *η*(*t*) from SFS summary data—is complicated by the fact that different population size histories can have identical expected sample frequency spectra. This problem, known as non-identifiability, has been extensively explored in the literature [22, 23, 24, 25, 26], and it is generally solved by preferring population size histories that have fewer changes and biologically unrealistic oscillations. Here, we use similar smoothing assumptions to treat this non-identifiability, as well as a compositional constraint that we explain next.

A new yet tractable identifiability problem arises in the MuSH inference setting. The effective population size *η*(*t*) and the mutation intensity *μ*(*t*) are mutually non-identifiable for all *t*, meaning that the expected SFS ****ξ**** is invariant under a modification of *η*(*t*) so long as a compensatory modification is made in *μ*(*t*). The non-identifiability of *η* and *μ* can be understood intuitively by example: an excess of variants of a given frequency can be explained by an historical population boom, which lengthens coalescent lines in the boom time interval, but it may be explained equally well by a period of increased mutation intensity with no demographic change.

While the overall mutation intensity is confounded with demography, the *composition* of the mutation spectrum—the relative mutation intensity of each mutation type—reveals itself in the *k*-SFS. This can also be understood intuitively: an excess of variants of a given frequency in only a single mutation type (one column of the *k*-SFS) cannot be explained by an historical population boom, because all mutation types are associated to the same demographic history. In this case, we would infer a period of increased relative mutation intensity for this mutation type. Because we cannot discern changes in total mutation rate, mushi assumes a constant total rate *μ*_0_, so that time variation in the rate of drift is modeled only in *η*(*t*). Figure 1c schematizes how we handle this constraint using a transformation technique from the field of compositional data analysis. Details are described in the Methods.

Even with this compositional constraint on the total mutation rate, many very different and erratic population histories may be equally consistent with an empirical *k*-SFS. As mentioned before, we overcome this by leveraging recently developed optimization methods to find smoothly regularized demographies and MuSHs. We penalize the model for three different types of irregularity. One penalty is familiar from the demographic inference literature: histories that feature rapid oscillations of the effective population size over time are disallowed in favor of similarly likely histories with effective population sizes that change less rapidly and less often. The second penalty may be more familiar to users of clustering methods such as STRUCTURE [29], where information criteria are used to favor explanations of the data with as few independently varying ancestry profiles as possible. Analogous to this, we favor models in which the mutation spectrum history matrix ****μ****(*t*) has low rank, meaning that there exist relatively few mutational signatures that independently vary in their intensity over time. The third regularization penalty is known as a classical ridge or Tikhonov penalty, favoring solutions with small *ℓ*_2_ norm, which speeds up convergence of the optimization without significantly affecting the solution. Figure 1d schematizes intuitions behind our regularization approach, and detailed formulation of our optimization problems and regularization strategies are in the Methods.

The intensity of all three regularizations can be tuned up or down by changing the values of user-specified hyperparameters. As the strength of regularization is increased, the method returns increasingly simple histories, but eventually this may result in a poor fit between the expected *k*-SFS and the empirical *k*-SFS. Users should tune the regularization parameters to seek histories that appear as simple as possible without over-smoothing, a process that is designed to be more straightforward than the parametric model specification that is required by many methods that infer demography from the SFS.

### Quantifying goodness of fit to the data

The likelihood of an empirical SFS given an expected SFS is often measured using a Poisson random field (PRF) approximation [30], which stipulates that, neglecting linkage, the observed number of sites with frequency *i* is Poisson-distributed around the expected number of sites of this frequency. This PRF approximation is easily generalizable to *k*-SFS data. Recall that **X** is the observed *k*-SFS matrix, so the SFS is **x***≡* **X1** (row sums over mutation types). In the Methods we show that the generalized PRF likelihood factorizes as ℙ(**X***| η, **μ***) = ℙ(**x***| η*) ℙ(**X***|* **x***, η, **μ***), with the first factor given by a Poisson and the second by a multinomial likelihood. We also show that the SFS **x** is a sufficient statistic for the demographic history *η* with respect to the *k*-SFS **X**. This means that estimation of *η* can be done by fitting the total SFS, which maximizes the first factor. Then the MuSH can be estimated by fitting the *k*-SFS, maximizing the second factor, conditioned on this *η* estimate.

## Results

### Reconstructing simulated histories

We first investigated the ability of mushi to recover histories in simulations where known histories are used to generate *k*-SFS data. Instead of simulating under the mushi forward model itself, we used msprime [31] to simulate a *succint tree sequence* describing the genealogy for 200 haplotypes of human chromosome 1 across all loci. This is a more difficult test, as it introduces linkage disequilibrium that violates our model assumptions. The results of this section can be reproduced with the supplementary notebook https://harrispopgen.github.io/mushi/notebooks/simulation.html

We used the human chromosome 1 model implemented in the stdpopsim package [32], which includes a realistic recombination map [33]. We used a difficult demography consisting of a series of exponential crashes and expansions, variously referred to as the “sawtooth”, “oscillating”, or “zigzag” history. This pathological history has been widely used to evaluate demographic inference methods [34, 35, 36, 28], and is available in the stdpopsim package as the Zigzag_1S14 model for use with msprime. Our simulated tree sequence contained about 250 thousand marginal trees.

We defined a MuSH with 96 mutation types, two of which are dynamic: one undergoing a pulse, and the other a monotonic increase. The total mutation rate varies due to these two components— introducing another model misspecification, since inference assumes only compositional changes. We placed mutations on the simulated tree sequence according to the historical intensity function for each mutation type, and computed the *k*-SFS.

Figure 2 depicts inference results for this simulation scenario. We find that mushi accurately recovers the difficult sawtooth demography for most of its history, but begins to over-smooth by the time of the third population bottleneck because little information survives in the SFS from this time period. The MuSH is accurately reconstructed as well, with both the pulse and ramp signatures recovered, and the remaining 94 components flat. The timing of the features in the MuSH also appears accurate, despite demographic misspecification that has the potential to distort the diffusion timescale.

**Figure 2:**
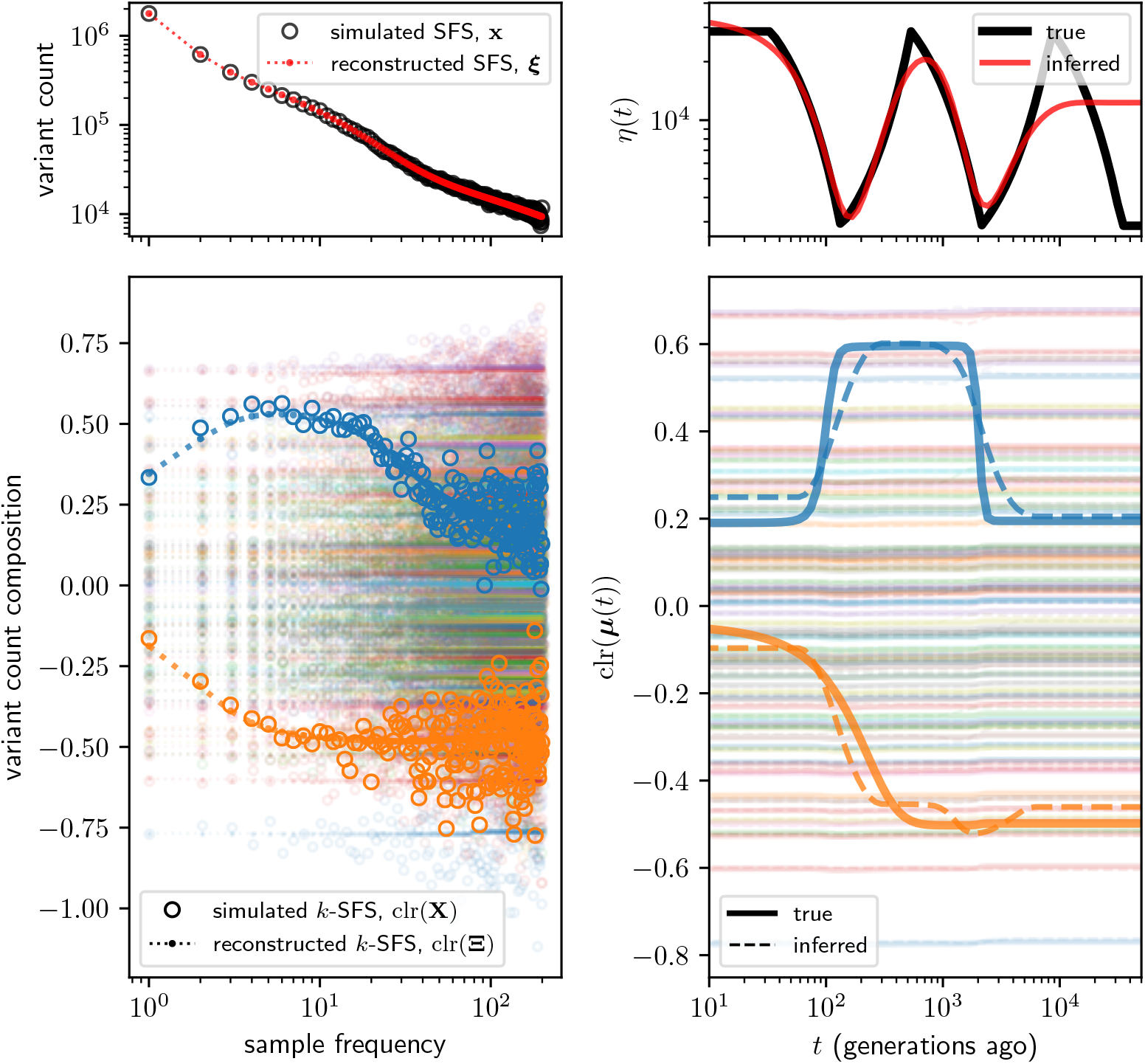
Simulation study of mushi performance. The sawtooth demography (top right) and a MuSH with 96 mutation types (bottom right, with two non-constant components in bold) were used to simulate 3-SFS data for *n* = 200 sampled haplotypes. The MuSH has a total mutation rate of about *μ*_0_ = 83, generating about 8.3 million segregating variants. The top left panel shows the SFS, and the bottom left shows the *k*-SFS as a composition among mutation types at each allele frequency (the two components corresponding to the non-constant mutation rates are in bold). Time was discretized with a logarithmic grid of 100 points.

One noteworthy feature of our fit to the sawtooth demography is the increasing tendency of mushi to smooth out older demographic oscillations without smoothing younger oscillations as aggressively. In contrast to methods such as the pairwise sequential Markov coalescent (PSMC) [37] that tend to infer large, runaway population sizes in the ancient past, mushi is designed such that the inferred history flattens in the limit of the ancient past. The same constraint underlies both PSMC’s ancient oscillations and our method’s ancient flattening: genomic data sampled from modern individuals cannot contain information about history older than the time to most recent common ancestor (TMRCA) of the sample, since mutations that occurred before then will be fixed, rather than segregating, in the sample. For example, we expect that population bottlenecks erase information about more ancient history, since they accelerate the fixation of variant sites that predate the bottleneck. While this information loss intuition holds for very general coalescent processes [38], the linearity in Theorem 1 enables us to make these statements precise for mutation rate history via spectral analysis of the operator 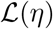. This is explored in detail for the case of a simple bottleneck demography in Appendix A.5, and the results may be reproduced from the supplementary notebook https://harrispopgen.github.io/mushi/notebooks/observability.html.

### Reconstructing the histories of human populations

With encouraging results from simulation experiments, we next set out to infer the histories of human populations from large publicly-available resequencing data. We computed a *k*-SFS for each of the 26 human populations from 5 continental ancestries sequenced in the 1000 Genomes Project (1KG) [27]. Our bioinformatic pipeline for computing the *k*-SFS for each 1KG population is detailed in the Methods, and a reusable implementation is provided in the mushi repository. Briefly, we augment autosomal biallelic SNPs in variant call data by adding triplet mutation type (*k* = 3) annotations, masking for strict callability and ancestral triplet context identifiability. Across 1KG populations the resulting number of segregating variants ranged from *~*8 million (population CDX) to *~*16 million (population ACB). We also computed the genomic target sizes for each ancestral triplet context, resulting in a total ascertained genome size of *~*2.0 Gb.

A few basic model parameters are defined as follows. We use a de novo mutation rate estimate of *μ*_0_ = 1.25 *×* 10^*−*8^ per site per generation [39], which corresponds to *~*25.4 mutations per *~*2.0Gb masked haploid genome per generation. For time calibration, we assume a generation time of 29 years [40]. To discretize the time axis, we use a logarithmically-spaced grid of 200 points, with the most recent at 1 generation ago, and the oldest at 200 thousand generations (5.8 million years) ago. Finally, we mask the last 10 entries in the SFS, which are more vulnerable to ancestral state mis-identification. Other details, including regularization parameter settings, are available in the supplementary notebook https://harrispopgen.github.io/mushi/notebooks/1KG.html, which reproduces the results of this section.

### Human demographic history

We used mushi to infer demographic history *η*(*t*) independently for each 1KG population. Figure 3 shows results grouped by super-population: African (AFR), Admixed American (AMR), East Asian (EAS), European (EUR), and South Asian (SAS). Broadly, we recover many previously-known features of human demographic history that are highly robust to regularization parameters, genomic masks, and SFS frequency masking: a *~*100 kya out-of-Africa bottleneck in non-Africans, a second contraction *~*10 kya due to a founder event in Finland (FIN), and recent exponential expansion of all populations. Histories ancestrally converge within each super-population, and super-populations converge at the most ancient times.

**Figure 3:**
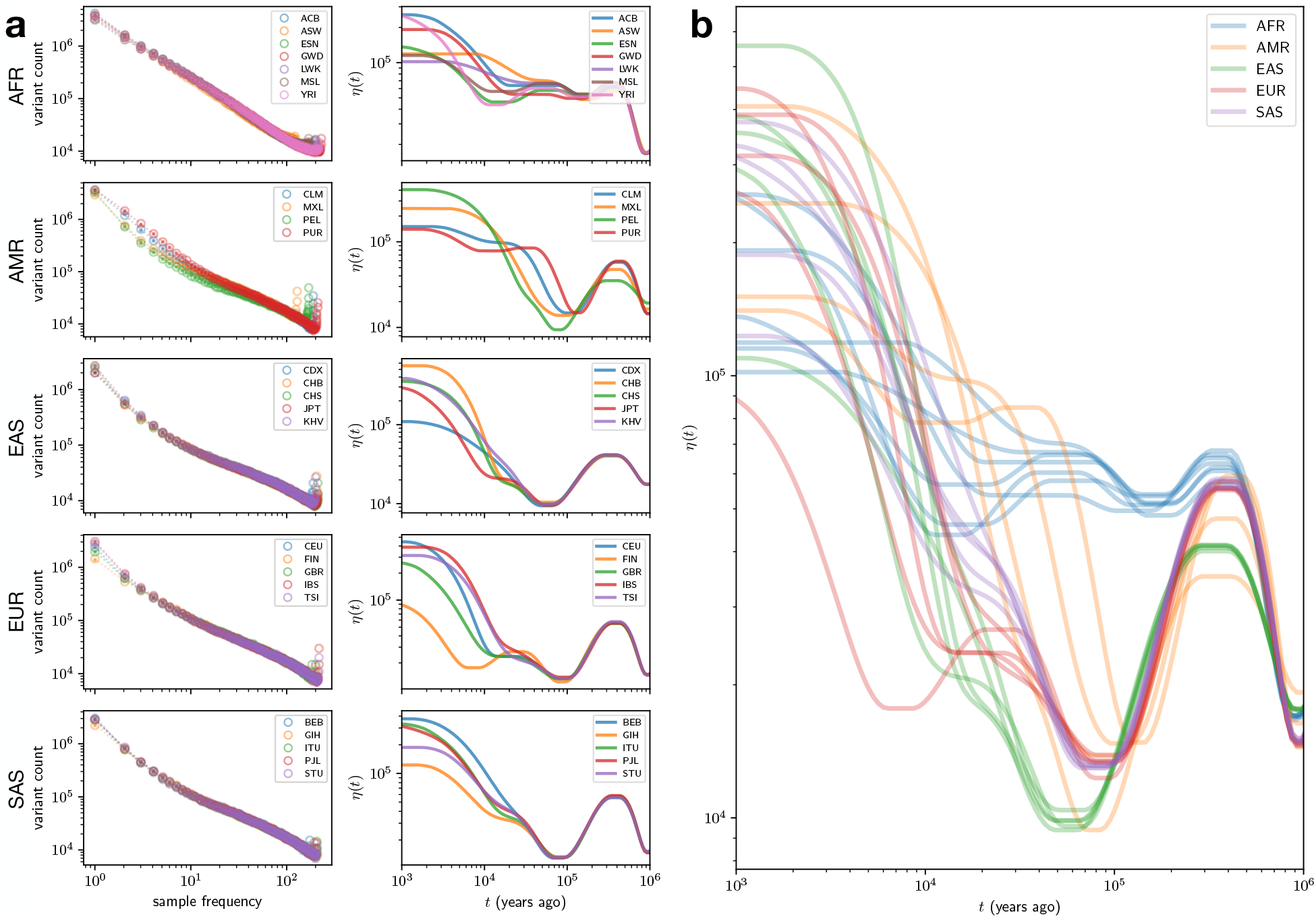
Effective population size histories for 1000 Genomes Project populations. **a.** The left column shows SFS data (open circles) for each population with separate panels for each super-population, as well as fits based on the expected SFS from the estimated demography history (points connected by dotted lines). The right column shows the corresponding demographic history *η*(*t*) estimates. **b.** The same *η*(*t*) estimates as in (b.) on common axes, to allow comparison of super-populations.

### Human mutation spectrum history

Each of our estimated demographic histories induces a mapping of population allele frequency onto a distribution of allele ages. With these distributions encoded in our model, we next used mushi to infer time-calibrated MuSHs for each population. First, to highlight the time calibration capabilities of mushi, we focus on the specific triplet mutation type TCC*→*TTC, which was previously reported to have undergone an ancient pulse of activity in the ancestors of Europeans, and is absent in East Asians [17, 18, 28]. To produce sharp estimates of the timing of this TCC pulse, we used regularization parameters that prefer histories with a minimum number of change points (see Methods). Figure 4a shows our fit to this component of the *k*-SFS for each EUR population, and Figure 4b shows the corresponding estimated component of the MuSH.

**Figure 4:**
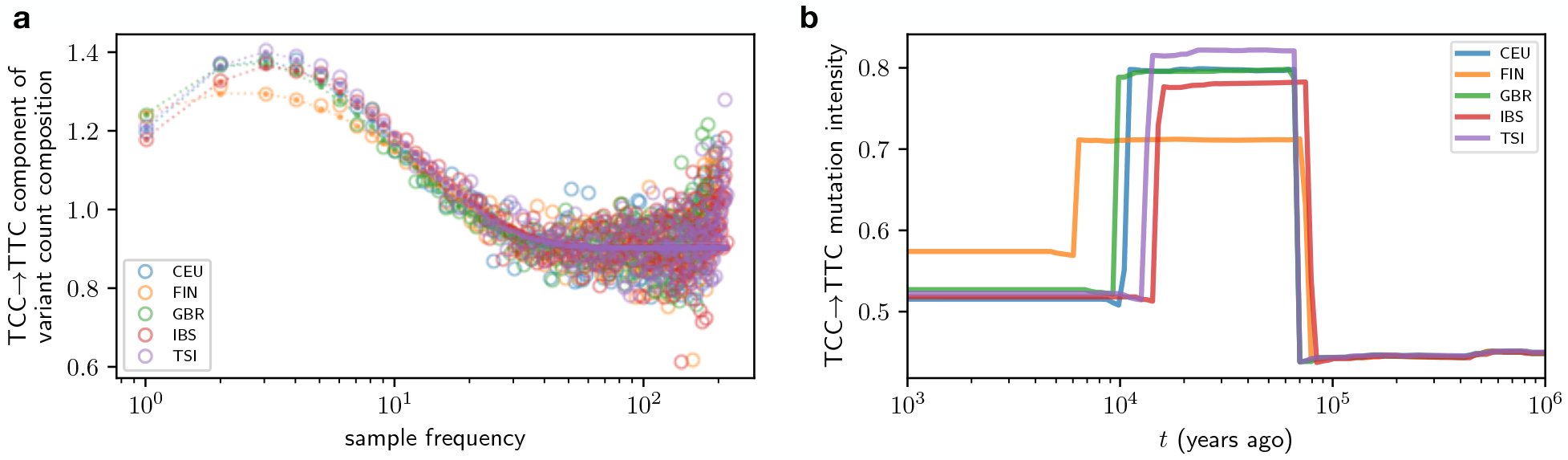
Timing of TCC *→* TTC pulse in Europeans. More accurate timing of a previously-reported pulse in TCC*→*TTC mutation rate in the ancestors of Europeans is enabled by joint inferencce of MuSH and demography. **a.** The relative composition of TCC*→*TTC variants in each frequency class for each EUR population (computed with the centered log ratio transform, see Methods), shows an excess at intermediate frequencies (open circles). The expectated values fit using the inferred MuSH are shown as points connected by dotted lines. **b.** The corresponding inferred TCC*→*TTC mutation intensity histories (in units of mutations per ascertained genome per generation).

With the consistent joint estimation performed by mushi, we find that the TCC pulse is much older than previously reported, beginning *~*80 kya. It is also possible to run mushi without estimating a new demographic history from the input data, but instead assuming a pre-specified demography. When we use the Tennessen, et al. history [41], which was assumed by Harris and Pritchard [18] in their estimate of the timing of the TCC pulse, we recover a pulse beginning around 15-20 kya, as previously estimated. We estimated a third set of European MuSHs conditioned on demographic histories that were inferred using the recently developed method Relate [28], which utilized the same 1KG input data that we analyze here, but leveraged linkage information as well as allele frequencies to infer population size changes. Conditioning on the Relate demographies also yielded younger estimates of the TCC pulse timing, but both pre-specified demographies fit the SFS poorly, indicating that demographic misspecification has likely distorted mushi’s time calibration (see section “TCC*→*TTC pulse in Europeans” of the supplementary notebook linked above). It is also likely that the mushi-inferred history fails to fit features of the data such as linkage disequilibrium patterns. If further advances in demographic inference manage to produce a history that fits both the SFS and orthogonal aspects of the data, this might necessitate further revisions to our best estimates of MuSH time calibration.

After our focused study of the TCC pulse, we aimed to more broadly characterize how human MuSH decomposes into principal mutation signatures varying through time in each population. We ran mushi on all 1KG populations using regularization parameters that favor smooth variation over time, rather than constraining the number of change points (see Methods). This resulted in an estimated MuSH for each population of the 26 populations in the 1KG data. Fits to the *k*-SFS and reconstructed MusHs are shown for each 1KG population in supplementary notebook section “Mutation spectrum histories for all populations”. We then normalized each MuSH by the genomic target size for each triplet mutation type, so that mutation rate is rendered site-wise, and stacked the population-wise MusHs to form an order 3 tensor. As pictured in Figure 5a, this tensor is a 3D numerical array with dimensions (num. populations) × (num. time points) × (num. mutation types) = 26 × 200 × 96. When we slice the array along the time axis, we obtain a series of matrices whose rows are the inferred mutation spectra of each 1KG population at a past time *t*. The numerical value of an entry in the tensor indicates the mutation rate (in units of mutations per site per generation) in a given population, at a given time, and for a given mutation type.

**Figure 5:**
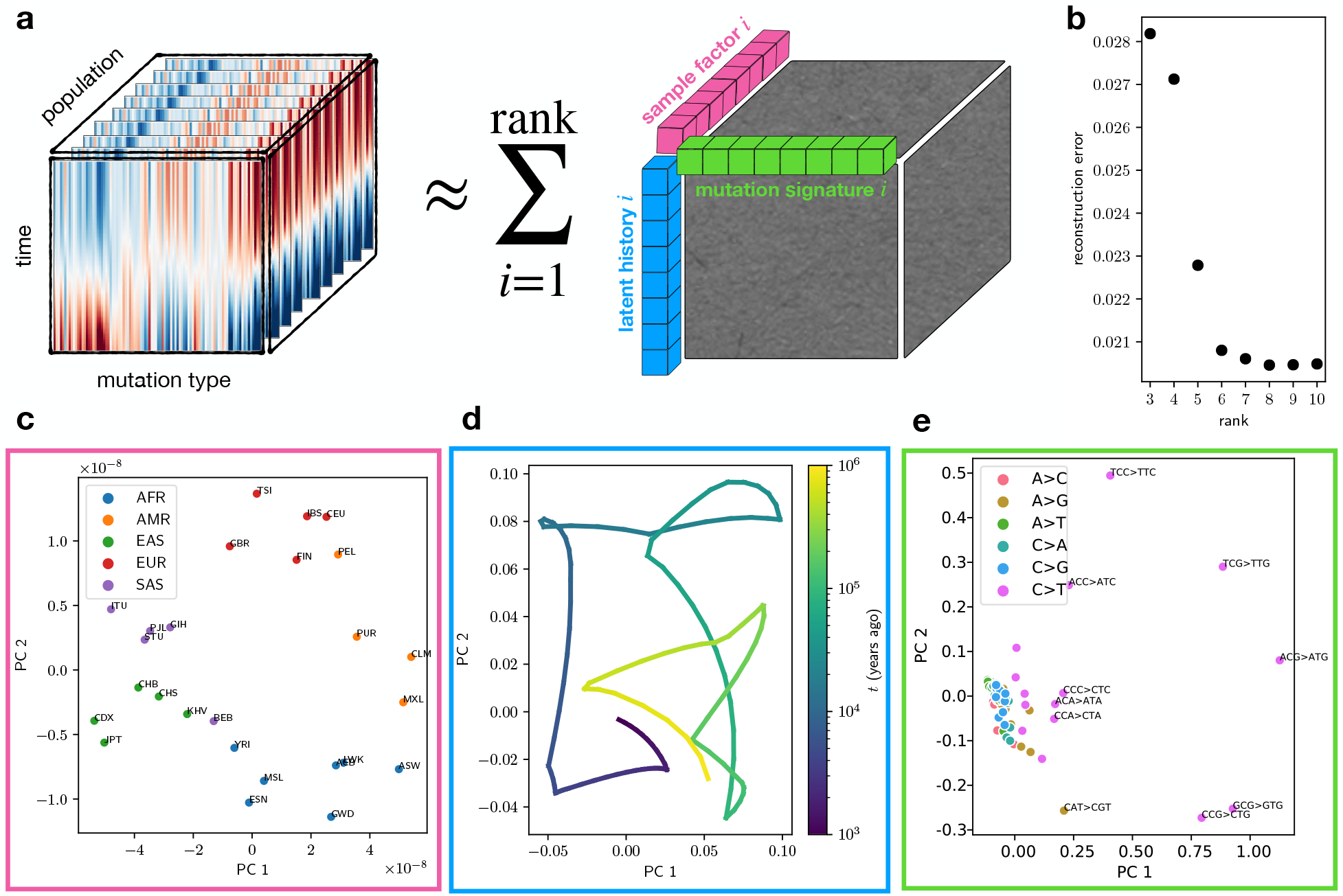
Decomposition of mutation spectrum histories for 1000 Genomes Project populations into nonnegative factors. **a.** Schematic of tensor decomposition, showing the MuSH for all populations stacked into a 3rd dimension, and approximated as the sum of tensors of rank 1. The set of rank 1 tensors in this sum are composed (via an outer product) of factors for populations, times, and mutation signatures, which are amenable to biological interpretation. **b.** Tensor reconstruction error over a range of ranks for NNCP decomposition, indicating rank 8 as a good approximation. **c.** 8-dimensional population factors projected to first 2 principal components. **d.** 8-dimensional time factors projected to first 2 principal components. **e.** 8-dimensional mutation signature factors projected to first 2 principal components. Overall, variation in the rates of select transitions account for most of the mutation spectrum variation between populations.

We used non-negative canonical polyadic tensor factorization (NNCP, reviewed by Kolda and Bader [42]) to extract factors in the population, time, and mutation type domains. This is analogous to extracting mutational signatures that form a low rank vocabulary for explaining the mutation spectrum variation between tumor mutational profiles. NNCP generalizes non-negative matrix factorization to tensors of arbitrary order. The addition of the time dimension means that each mutational signature is associated with a dosage that can jointly increase or decrease over the histories of all populations.

Briefly, we hypothesize that the MuSH tensor can be approximated by a sum of a few rank-1 tensors (Figure 5a). This is tantamount to assuming that most evolving mutational processes are shared across multiple populations, possibly with different relative intensities over time. We find that a tensor of rank 8, which describes a set of 8 mutational processes, can accurately represent the 1KG MuSH tensor (Figure 5b). This NNCP decomposition results in 26 × 8, 200 × 8, and 96 × 8 factor matrices for population, time, and mutation type, respectively. Figure 5c–e projects each set of factors from 8 dimensions to 2 principal components for visualization. The population factors (Figure 5c) clearly cluster by super-population. The time factors (Figure 5d) trace out a continuous trajectory in factor space for the set of all populations, which is expected since regularization in mushi imposed smoothness in the time domain. The mutation type factors (Figure 5e) show a number of mutation types with distinct outlier behavior, including TCC*→*TTC, as expected.

We next recast the MuSH for each population in terms of the 8 mutation signatures that comprise the tensor factors, capturing covariation among the set of 96 triplet mutation types with the smaller set of signatures. This allows us to characterize and biologically interpret the time dynamics of each mutation signature in each population. Figure 6a shows the 8 mutation signatures as loadings in each triplet mutation type. Figure 6b shows how each of these 8 signatures varies through time in each 1KG population (computed by projecting 96-dimensional spectra to the 8 mutational signatures in each population at each time). Signature 3 fits the profile of the TCC pulse that affects Europeans, South Asians, and European-admixed Amerindians, containing all previously reported minor components of the pulse such as ACC*→*ATC and CCC*→*CTC. Signatures 1 and 5, which are consistent with deamination of CpG sites, are consistently enriched in rare (young) variants across populations, which is likely due to a combination of purifying selection and biased gene conversion. Biased gene conversion disfavors the increase in frequency of C/G*→*A/T mutations (also called strong-to-weak mutations), and many CpG sites are conserved due to their role in the regulating chromatin accessibility as well as gene expression. Signatures 2 and 6 are enriched for common (old) variants, and have high loadings of A*→*G, which is consistent with the action of biased gene conversion to select for the retention of weak-to-strong mutations.

**Figure 6:**
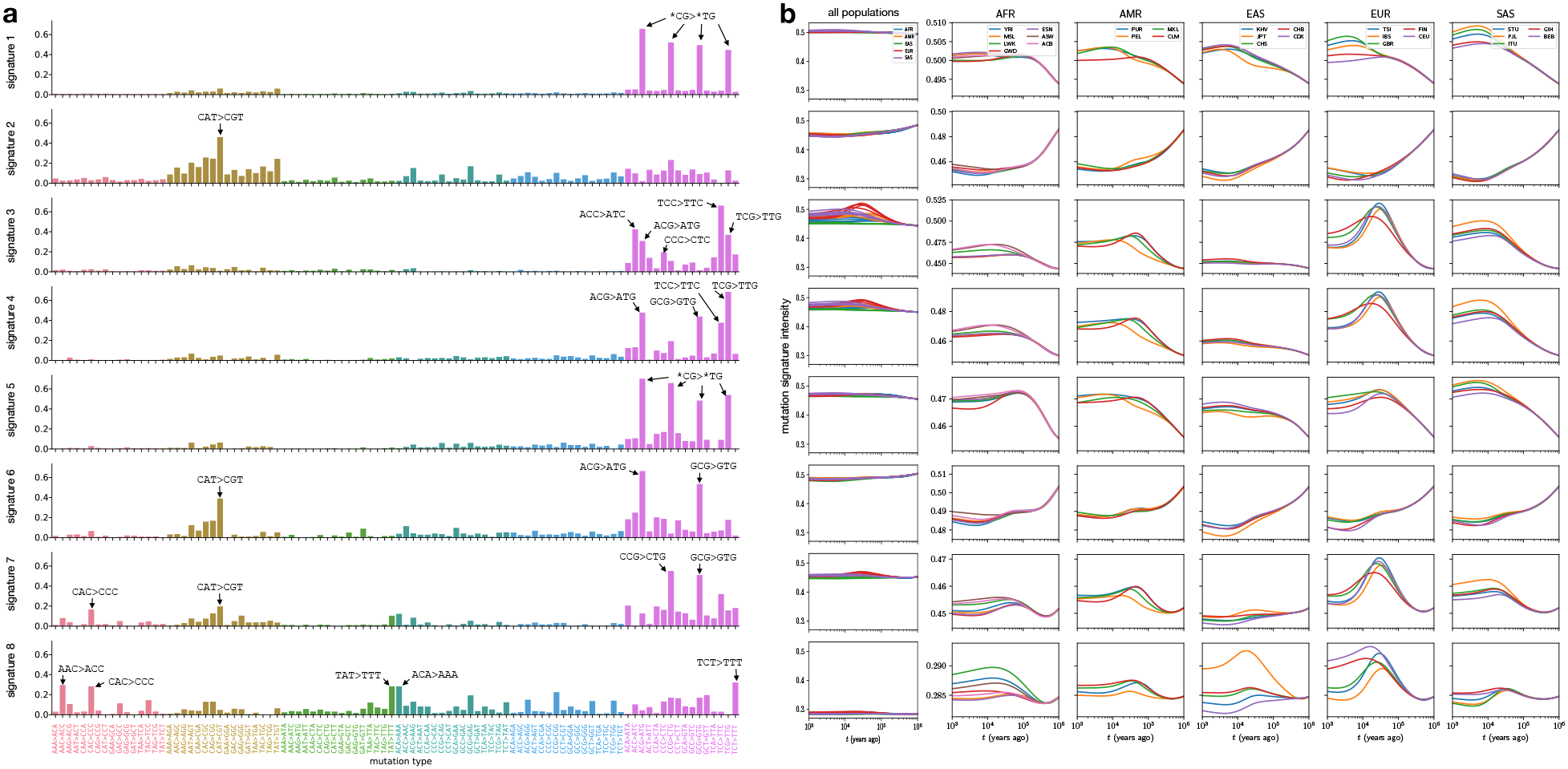
Dynamics of mutation signatures in the history of 1000 Genomes Project populations. **a.** Triplet mutation signatures, shown as loading into triplet mutation types for each signature (rows) **b.** Historical dynamics of each mutation signature in each 1KG population, with rows corresponding to signatures in (a). The first column shows all populations on common axis ranges to indicate relative scale, and the remaining columns show the same histories for each super-population, with ranges for each signature.

Although the time profiles of these 8 signatures appear to be modulated by biased gene conversion, they also vary between populations at recent times and cannot be explained by a selective force acting uniformly on all non-GC-conservative mutations. Signature 8 fits the profile of a signature reported to be enriched specifically in the Japanese population [18]; though this signature may stem from a subtle cell line artifact affecting the Japanese Hap Map samples [43], it is still a feature of the 1KG data that is expected to fit the profile of a population-specific mutational signature. Signature 4, which is dominated by C*→*T transitions, is enriched in Europeans and South Asians relative to East Asians and Africans, charting the time course of a trend that was previously reported in empirical heat map data [18]. Another reported trend is the existence of differences between populations in the rates of CA\**→*CG* mutations which can be explained by differences between populations in the recent dosages of signatures 7 and 8.

Finally, we used uniform manifold approximation and projection (UMAP) [44] to compute a 2D embedding of mutation *signature* histories (after initially decomposing the MuSHs into 8 mutation signatures as described) of each 1KG population at each time point. Figure 7a shows this embedding with all times in the same axes. Despite performing independent inferences for each population’s MuSH, we see recapitulation of trees of population and super-population ancestry. Figure 7b shows the same embedding with the time coordinate resolved as a 3rd dimension.

**Figure 7:**
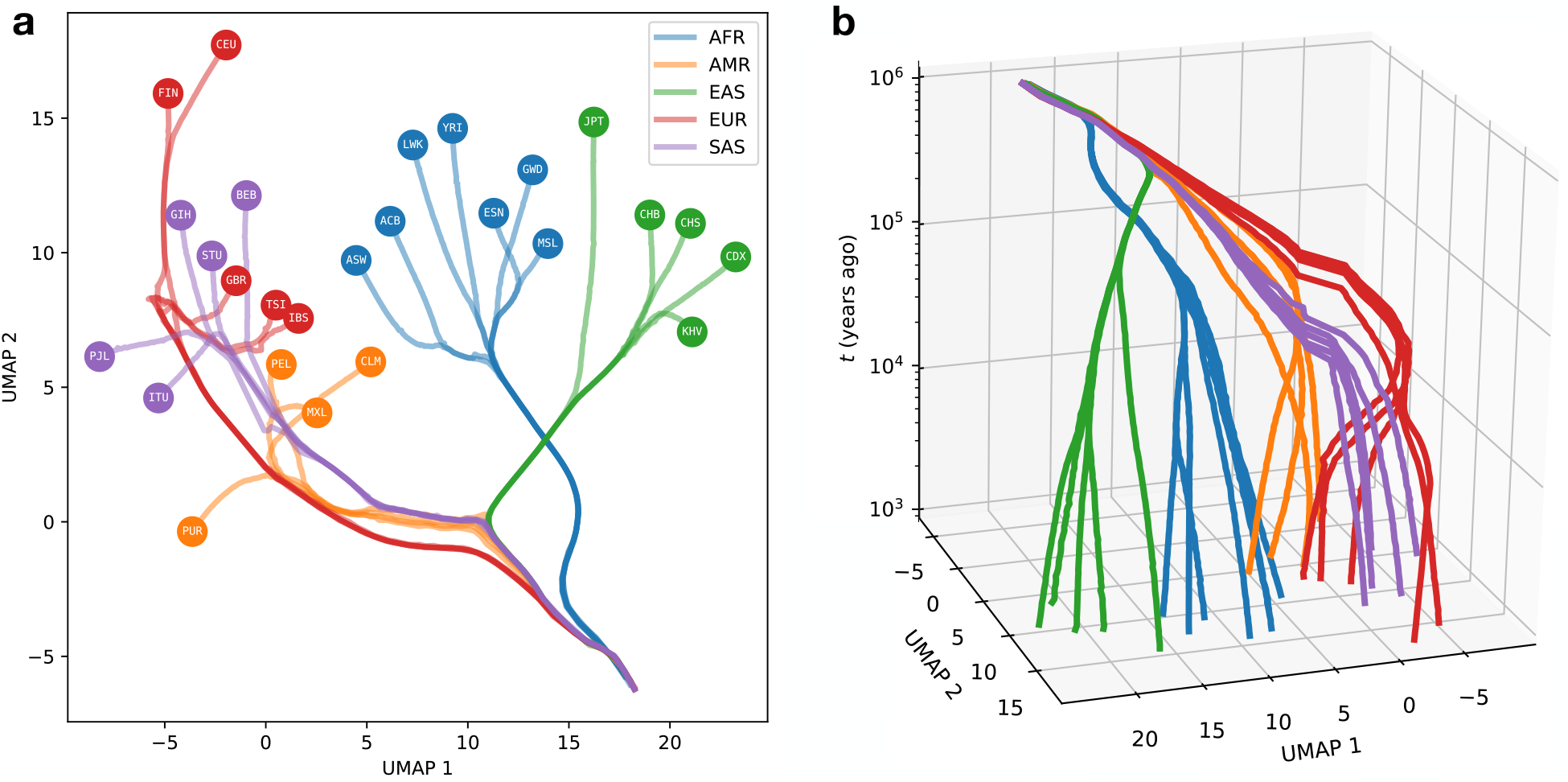
Global divergence in the mutation signature history of 1000 Genomes Project populations. **a.** UMAP embedding of mutation signature histories was initialized using the first two PCs of the time-domain factors, and then performed with default parameters. **b.** Equivalent embedding with time coordinate added as a 3rd dimension.

## Discussion

It is becoming increasingly clear that mutation spectrum variation is a common feature of large genomic datasets, having been discovered and formally reported in population sequencing panels from humans, great apes, and mice [18, 45, 46]. Initial reports on the existence of such variation were mostly qualitative in nature, focused on enumerating which populations exhibit robust variation along this newly characterized dimension and putting bounds on the possible contributions of bioinformatic error. Here, we have introduced a novel quantitative framework for characterizing mutation spectrum evolution within populations, which utilizes variation of all ages from unphased whole genome data to resolve a time-varying portrait of germline mutagenesis. Our method mushi can decompose context-augmented sample frequency spectra into time-varying mutational signatures, regardless of whether those signatures are sparse and obvious like the European TCC pulse or represent more subtle concerted perturbations of mutation rates in many sequence contexts. Previous estimates of the timescale of mutation spectrum change were restricted to sparse signatures that are more obvious but less ubiquitous than diffuse signatures appear to be [18, 28].

Not all of the temporal structure unveiled by mushi can be interpreted as time variation in the germline mutational processes. Some time variation in signature dosage is consistent with the action of biased gene conversion, and there is no automated mechanism to flag signatures that have suspicious hallmarks of cell line artifacts [43]. The strengths of mushi are to automate the visualization of deviations from mutation spectrum uniformity and quickly localize them to particular populations, frequency ranges, and time periods, enabling straightforward scrutiny and the design of downstream investigations of their validity and ontogeny.

Although mushi’s most novel feature is the ability to infer mutation spectrum variation over time, it includes a demographic inference subroutine with several advantages over existing demographic inference methods. Ours is only the second method to infer population size changes non-parametrically from SFS data [47], and its state-of-the-art regularization methods yield population size histories with some more desirable properties than other methods for non-parametric effective population size history inference. With mushi, adaptation to temporally localized smoothness levels is much better than with smoothing splines [48]. Histories inferred by mushi stabilize to a constant size in the limit of the ancient past rather than exhibiting runaway behavior due to overfitting, and the use of sample allele frequencies rather than phased whole genomes should make the method broadly useful to researchers working on non-model organisms, which are still beyond the scope of many state-of-the-art methods that require long sequence scaffolds and phased data. The software is also very fast, returning results in seconds on a modest computer, and is designed to be easily used by biologists familiar with scripting in Python.

The mushi model calibrates the times at which mutational signatures wax and wane using a demographic model inferred from the same input allele frequency data from which the signatures themselves are extracted. However, it can also calibrate its timescale using a user-specified demographic history, which reveals that the timing of transient events like the TCC pulse in Europe are exquisitely sensitive to underlying assumptions about effective population size. When we input demographic histories previously inferred from other datasets, we conclude that the TCC pulse began 15,000 to 30,000 years ago, comfortably later than Europeans’ divergence from East Asians, which were not affected by the TCC pulse. However, inferred demographic histories are notoriously poor at predicting the distributions of genomic summary statistics other than the ones that were used to fit the models [49], and these external demographic history estimates yield poor fits to the 1KG SFS data. When we use mushi to estimate population histories that do fit the 1KG sample frequency spectra well, we estimated a surprisingly old start time to the TCC pulse, around 80 kya, which is older than any estimates of European/East Asian divergence times. This might invoke ancient population structure to explain the allele frequency distribution of excess TCC*→*TTC mutations in Europe. For example, the pulse may have initially been active in a basal European population that diverged from East Asians earlier than other populations that contributed to modern European ancestry. This puzzle points to the need for future work modeling mutation spectrum evolution jointly with more complex demographic history involving substructure and migration between populations. It also points to the tantalizing possibility that the distribution of mutational signatures could provide extra information about hard-to-resolve substructure and gene flow between populations that no longer exist in “pure” form today.

Although powerful new methods for inferring ancestral recombination graphs (ARGs) ultimately have the potential to estimate more accurate demographic histories than can be accomplished by fitting more compressed SFS data, these methods are still in a relatively early stage of development. In the method Relate [28], mutation rate history is approximately inferred from an ARG using independent marginal estimates for each epoch in a piecewise-constant history. This avoids joint inference over all epochs—which can also be formulated as a linear inverse problem—by ignoring mutation rate variation within branches. Although this lowers computational complexity, it comes at the cost of estimator bias that is not well characterized.

Our results underscore the importance of using more compressed summary statistics to validate inference results. In theory, an ARG contains perfect information about the SFS as well as additional information about linkage, meaning that demographic history inferred from an ARG should be consistent with the SFS. The differences between our SFS-inferred histories and Relate-inferred histories have significant implications with regards to the joint distribution of allele age and allele frequency. This could affect claims about the timing of gene flow and selection in addition to the claims about the timing of the TCC pulse that we focus on in this paper. Until the field of demographic inference achieves its holy grail of inferring histories that are compatible with all features of modern datasets, it will be important for researchers to practice inferring histories from different data summaries including classical, compressed statistics like the SFS in order to understand the sensitivity of various biological and historical claims to methodological eccentricities.

## Acknowledgements

WSD thanks the following individuals for discussions and feedback that greatly improved this work: Peter Ralph, Andy Kern, and members of the Kern-Ralph colab; Jeff Spence; Stilianos Louca and Matthew Pennell; Joe Felsenstein; Leo Speidel; Matthias Steinrücken; Andy Magee; Sarah Hilton; Erick Matsen and members of the Matsen group; UW Popgenlunch attendees Elizabeth Thompson, Phil Green, Mary Kuhner; members of the Harris lab. KDH thanks Aleksandr Aravkin for suggesting proximal splitting methods and for other discussions. WSD was supported by the National Institute Of Allergy And Infectious Diseases (F31AI150163) and by the National Human Genome Research Institute (T32HG000035-23) of the National Institutes of Health. KDH was supported by a Washington Research Foundation Postdoctoral Fellowship. KH was supported by the National Institute of General Medical Sciences (1R35GM133428-01) of the National Institutes of Health, a Burroughs Wellcome Career Award at the Scientific Interface, a Pew Biomedical Scholarship, a Searle Scholarship, and a Sloan Research Fellowship.

## Author contributions

Initial conception and formal analysis was done by WSD. WSD and KDH developed computational methods and software implementation. WSD performed data analysis in consultation with KDH and KH. The manuscript was initially drafted by WSD, and the all authors contributed to the final draft.

## Methods

### The expected SFS is a linear transform of the mutation intensity history

We work in the setting of Kingman’s coalescent [50, 51, 52, 53], with all the usual niceties: neutrality, infinite sites, linkage equilibrium, and panmixia [54, 55]. In Appendix A we retrace the derivation by Griffiths and Tavaré [56] of the frequency distribution of a derived allele conditioned on the demographic history, while generalizing to a time inhomogeneous mutation process. We make use of the results of Polanski et al. [57, 58] to facilitate computation. We use the time discretization of Rosen et al. [26], and adopt their notation. Detailed proofs can be found in the Appendix.

With *n* denoting the number of sampled haplotypes, denote the expected SFS column vector ****ξ**** = [*ξ*_1_ *… ξ_n−_*_1_]^T^, where *ξ_i_* is the expected number of variants segregating in *i* out of *n* haplotypes. Let *η*(*t*) denote the haploid effective population size history, with time measured retrospectively from the present in Wright-Fisher generations. Note that *η*(*t*) = 2*N* (*t*) for diploid populations. Let *μ*(*t*) denote the mutation intensity history, in units of mutations per ascertained genome per generation, understood to apply uniformly across individuals in the population at any given time. Under these model assumptions, we obtain the following theorem, whose detailed proof can be found in Appendix A.1.

#### Theorem 1.

*Fix the number of sampled haplotypes n. Then, for all bounded functions η* : ℝ_≥0_ → ℝ_*>*0_ *and μ* : ℝ_*≥*0_ *→* ℝ_*≥*0_, *the expected SFS is* 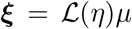, *where* 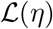 *is a finite-rank bounded linear operator parameterized by η that maps mutation intensity histories μ to* (*n −* 1)*-dimensional SFS vectors **ξ**. Viewed as a nonlinear operator on η*, 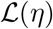 *is also bounded. In particular, 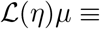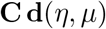, where* **C***is an* (*n −* 1) × (*n −* 1) *constant matrix with elements that can be computed recursively, and* **d**(*η, μ*) *is an* (*n −* 1)*-vector with elements*

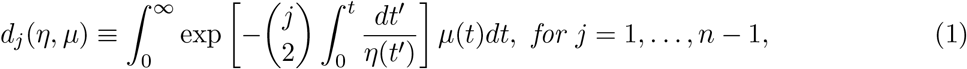

*which is linear in μ and nonlinear in η.*

Recursions for computing **C** can be procedurally generated using Zeilberger’s algorithm [59], which we detail in Appendix A.2).

In order to partition the expected SFS ****ξ**** by *k*-mer mutation type, we promote the (*n −* 1)-element expected SFS vector ***ξ*** to the (*n −* 1) × *κ* expected *k*-SFS matrix **Ξ** (not to be confused with the simultaneous multiple merger coalescent of Schweinsberg [60, 38] or the “SFS manifold” of Rosen et al. [26]). Similarly, the mutation intensity history function *μ*(*t*) is promoted to the *κ*-element mutation spectrum history ****μ****(*t*), a column vector with each element giving the mutation intensity history function for one mutation type. Then Theorem 1 generalizes to

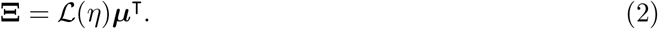

As in Theorem 1, the time coordinate is integrated over by the action of the operator 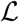.

We use the notation **X** to denote a sampled *k*-SFS matrix, i.e. the (*n −* 1) *× κ* matrix containing the sample counts for each mutation type. By construction, **Ξ***≡* 𝔼[**X**].

### Compositional modeling leads to identifiable mutation spectrum histories

As mentioned in the summary methods, the effective population size *η*(*t*) and the mutation intensity *μ*(*t*) are non-identifiable for all *t*, meaning that the expected SFS ****ξ**** is invariant under a modification of *η* so long as a compensatory modification is made in *μ*. We now demonstrate this formally by introducing a change of variables that measures time in expected number of coalescent events since the present, i.e. the diffusion timescale [22, 26]. Let 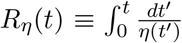, and substitute *τ ≡ R_η_*(*t*) in (1) to give

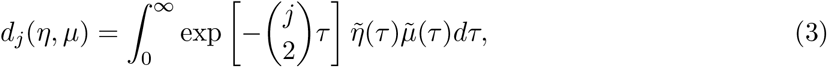

where 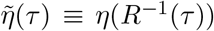 and 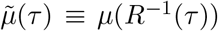. In this timescale, we see *η* and *μ* appear as a product on the right of (3). This means we cannot jointly infer *η* and *μ*, since only their product influences the data. This non-identifiability is similarly manifest by a change of variables to measure time in expected number of mutations.

Because we cannot discern changes in total mutation rate, we assume a constant total rate *μ*_0_, so that time variation in the rate of drift is modeled only in *η*(*t*). A MuSH with *κ* mutation types can then be written as ***μ***(*t*) = *μ*_0_***ν***(*t*), where 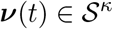 for all *t*, and 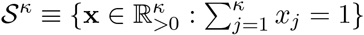 denotes the standard simplex. We call the relative mutation spectrum history ****ν****(*t*) a *composition*, and employ techniques from compositional data analysis [62, 63].

To avoid difficulties arising from optimizing directly over the simplex, we represent compositions using Aitchison geometry [62]. Briefly, analogs of vector-vector addition, scalar-vector multiplication, and an inner product are defined for compositions, and the simplex is closed under these operations. It is then possible to construct an orthonormal basis in the simplex ****Ψ****_1_*,…, ***Ψ***_k−_*_1_ using the Gram-Schmidt orthogonalization. We first introduce the *centered log ratio transform* of some 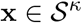, defined as

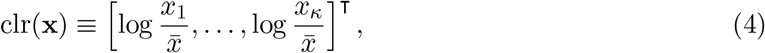

where 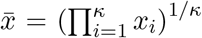 denotes the geometric mean. The inverse transform clr^*−*1^ is the softmax function.

The *isometric log ratio transform* and its inverse allow us to transform back and forth between the simplex and a Euclidean space in which we will cast our optimization problem. The transform ilr : 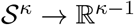 and its inverse are defined as

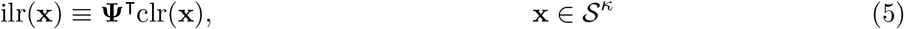

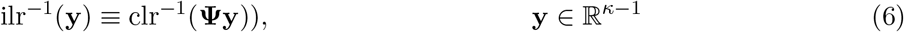

where **Ψ** ≡ [***Ψ***_1_ … ***Ψ***_*κ*−1_] is the *κ ×* (*κ −* 1) matrix of basis vectors. To build intuition about this transformation, which is an isometric isomorphism, we highlight the following behaviors: First, the center of the simplex maps to the origin in the Euclidean space. Second, approaching a corner of the simplex, i.e. with a component of the composition vanishing, corresponds to diverging to infinity in some direction the Euclidean space. Finally, a ball in the Euclidean space maps to a convex region in the simplex that is more distorted the further the ball is from the origin. These intuitions are illustrated in Figure 1c.

We use the convention that the clr and ilr act row-wise on matrices. Finally, we introduce the ilr-transformed MuSH: **z**(*t*) *≡* ilr(****μ****(*t*)) and write (2) as

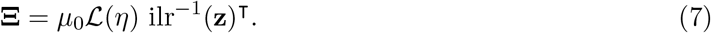

Again, the time coordinate is integrated over by the action of the linear operator. Although the forward model is non-linear in **z**(*t*), it is convex given the convexity of the softmax function that appears in ilr^*−*1^(*·*).

### Formulating and solving the inverse problem for population history given genomic variation data

The inverse problem (8) is ill-posed in general, meaning many very different and erratic histories can be equally consistent with the data [64]. We deal with this problem using regularization, seeking solutions that are constrained in their complexity without sacrificing data fit. We leverage recently-developed optimization algorithms to find regularized demographies and MuSHs.

### Time discretization

For numerical implementation, we need finite-dimensional representations of *η*(*t*) and **z**(*t*). We use piecewise constant functions of time on *m* segments [*t*_0_*, t*_1_), [*t*_1_*, t*_2_)*,…,* [*t_m−_*_1_*, t_m_*) where the grid 0 = *t*_0_ *< t*_1_ *< · · · < t_m−_*_1_ *< t_m_* = *∞* is common to *η*(*t*) and **z**(*t*). We take the boundaries of the segments as fixed parameters and, in practice, use a logarithmically-spaced dense grid of hundreds of segments to approximate infinite-dimensional histories. Let the *m*-vector **y** = [*y*_1_*,…, y_m_*]^T^ denote the population size *η*(*t*) during each segment, and define the *m ×* (*κ −* 1) matrix **Z** as the constant ilr-transformed MuSH **z**(*t*) during each segment. In Appendix A.3, we show that equation (7) discretizes to the following matrix equation

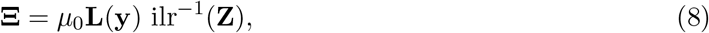

where the (*n*− 1) × *m* matrix **L**(**y**) is fixed given a fixed demographic history **y**. The transformation ilr^*−*1^(**Z**) is applied to each time point, i.e. row of **Z**, independently.

### Regularization

We implement three different regularization criteria: smoothness of the solutions **y** and **Z** (hypothesizing that the time variation of *η*(*t*) and **z**(*t*) is not excessively erratic), limited complexity of the matrix **Z**(hypothesizing that the number of independently evolving mutational signatures is much less than the number *κ* of distinct mutation types), and improved numerical conditioning of the problem. These goals are in some cases overlapping, but we add a regularization term for each one. Before computing the penalties on the demography **y**, we apply a log transform, because variation over orders of magnitude is expected from population crashes and exponential expansions. This also has the benefit of enforcing non-negative solutions. We now explain the regularizations in detail.

Our first regularization encourages smoothness in the time domain, as well as a limited number of change points, preferring to fuse consecutive segments of the history to the same value. This can be achieved by penalizing *ℓ*_1_ or *ℓ*_2_ norms of the time derivatives of log *η*(*t*) and **z**(*t*). In the discrete setting, the derivative operator can be approximated by a matrix **Δ** of first differences. This leads to the smoothing penalties 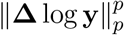 and 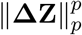. The penalty with *p* = 1 constrains the total number of time points at which a change in the function occurs and is referred to as a fused LASSO or total variation (TV) penalty [65, 66, 67]. Using *p* = 2 is called a spline penalty, as it enforces 1st-order smoothness analogous to differentiability [68]. Many demographic inference methods fit models composed of a small number of constant or exponential epochs that are motivated by prior knowledge about population histories. Although our histories are represented on a dense time grid, our regularization fuses neighboring time points to discover longer epochs of constant or smoothly varying behavior, while remaining flexible to capture more complicated behavior if the data justify it.

Second, because specific mutation processes may affect multiple mutation types, it is reasonable to assume that a small number of latent processes drive the majority of the variation across mutation types. We thus hypothesize that **Z** can be approximated by a low-rank matrix and propose two regularizations to enforce this. Let ***σ*** be the vector of singular values of **Z***−* **Z**_ref_, where **Z**_ref_ is a reference, or baseline, MuSH taken to be the MLE constant solution by default. We use the nuclear norm ||**Z***−* **Z**_ref_|| _*\**_ = ****||σ||**** _1_ as a *soft* rank penalty, as it is the convex envelope of the rank function [69]. The soft rank penalty constrains the number of non-zero singular values, while also shrinking them toward zero. As an alternative to the soft rank penalty we also implement a *hard* rank penalty, which directly penalizes rank(**Z***−* **Z**_ref_) = ***||σ||*** _0_, equal to the number of nonzero singular values. The hard rank penalty results in a singular value thresholding step without shrinkage in the resulting algorithm, and it is not convex. Either of these rank regularizations assure that **Z** is a low-rank perturbation of the constant solution **Z**_ref_. Although the MuSH represents the history of each of *κ* mutation types, this attempts to explain them using a smaller set of mutation signatures.

Finally, we include classical *ℓ*_2_ (also called ridge or Tikhonov) penalties on both log **y** and **Z**. A small amount of this kind of regularization speeds up convergence without significantly influencing the solution. For the ridge penalty on the demography **y**, we use a generalized Tikhonov term 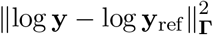 that allows the option of shrinking toward a reference demography **y**_ref_. Here **Γ** is a positive definite weight matrix which can be used to vary the strength of shrinkage across the time domain, and the notation 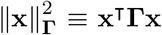 denotes the weighted norm squared. Note that the smoothing spline penalty is also of this form, but with the indefinite matrix **Δ**. By default we use the MLE constant history for **y**_ref_, and **Γ**= **I** (the identity matrix) to speed the convergence of the **y** problem. Similarly, the ridge penalty on the MuSH is a generalized Tikhonov term for each mutation type 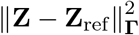, where the notation 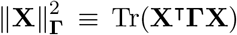 denotes the square of the weighted Frobenius norm. Although we model each population independently from the others, the generalized Tikhonov penalty can also be used to fuse the histories of populations that are known to share ancestry. For inferring 1KG demographies, we first performed inference for the YRI population using the default constant **y**_ref_ and **Γ**= **I**. For the other populations, we use the YRI history for **y**_ref_, and a diagonal 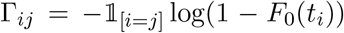, where *F*_0_ is the CDF of the TMCRA of the focal population using a constant demography estimate. This applies essentially no shrinkage for most of the history, but ramps up shrinkage toward YRI at times pre-dating the focal population’s TMRCA.

**Likelihood factorization: The SFS is a sufficient statistic for the demographic history with respect to the *κ*-SFS**

The PRF neglects linkage disequilibrium to model the probability of the SFS **x** given the expected **SFS *ξ*** as independent Poisson random variables for each sample frequency

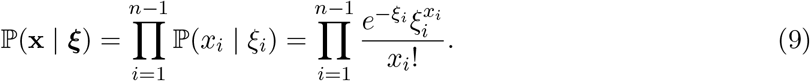

We similarly model the *k*-SFS as generated by independent mutational targets for each mutation type.

#### Proposition 1.

*The standard PRF indexed on sample frequencies generalizes to be indexed on the 2D grid of sample frequency and mutation type, and factorizes as* 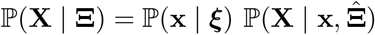, with 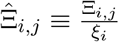. Here, ℙ(**x***| **ξ***) *is the Poisson distribution* (9)*, and* 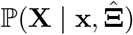 *is multinomial.*

*Proof.* We have that

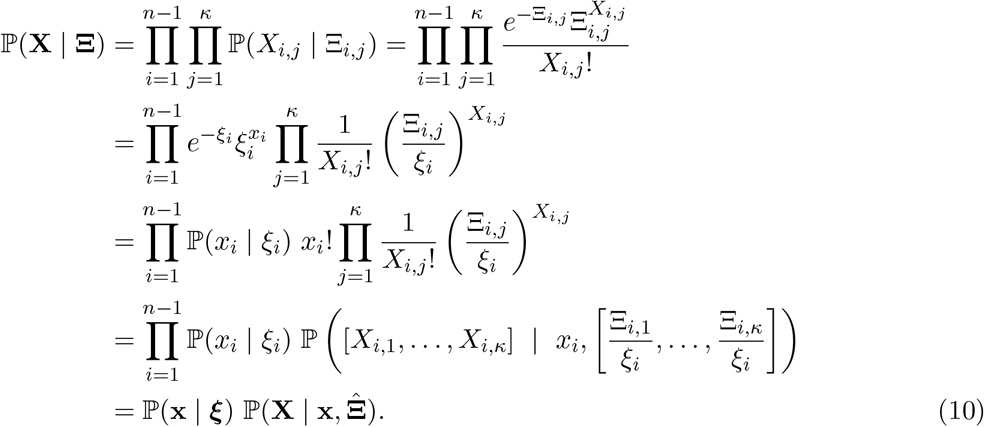

In the last two lines we’ve recognized the multinomially distributed mutation type partitioning of counts in each sample frequency *i*, with the rows of 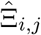 defining a multinomial parameter vector for each sample frequency *i*. The factorization of independent Poissons into an aggregate Poisson and a multinomial is a well-known result often called “Poissonization” [70].

Next we restore the *η* and ***μ*** dependence of ***ξ*** and **Ξ** (with fixed total mutation rate *μ*_0_) so (10) gives the factorization in the main text

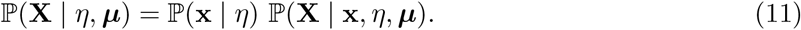

#### Lemma 1.

*If the total mutation rate is a constant **μ***(*t*) = *μ*_0_ *∈* ℝ_*>*0_, *then the SFS* **x** *is a sufficient statistic for η with respect to the k-SFS* **X**.

Lemma 1 is proved via a Poisson thinning argument in Appendix A.4. The result is intuitively obvious because information about historical coalescence rates recorded in the SFS does not change if we further specify how mutation counts are partitioned into different mutation types; this only adds information about relative mutation rates for alleles with a given age distribution. Although *η* appears in the second factor of (11), this only serves to map the MuSH rendered on the natural diffusion timescale 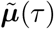 to time measured in Wright-Fisher generations. Because this map is one-to-one, there is no statistical information about *η* in **X** not already present in **x**. That is, 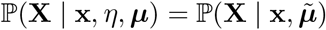.

This sufficiency is important from an inference perspective, because it means we can sequentially infer demography from the SFS, then infer the MuSH from the *k*-SFS with the demography fixed from the previous step. Sufficiency implies that the negative log-likelihood factors into the sum of two losses. We thus formulate two sequential optimization problems using negative log-likelihoods from the factors (11) as loss functions for assessing data fit. Recall that **y** and **Z** are the discrete forms of *η* and ***μ***, respectively, **Ξ** is given by equation (8), and ***ξ*** is given by the row sums of **Ξ** and thus independent of **Z**. Neglecting constant terms, the two loss functions are

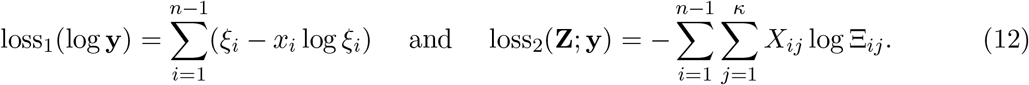

As with regularization, we parameterize in terms of log **y**.

### Optimization problems for mushi

We infer demography and MuSH by minimizing cost functions that combine the loss functions above, which measure error in fitting the data, with regularization. This may be considered a penalized likelihood method and, from a Bayesian perspective, may be interpreted as introducing a prior distribution over histories. Inference of log **y** and **Z** is performed sequentially. We first initialize log **y**= **y**_ref_ using the elementary formula for the MLE constant demography 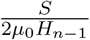 where 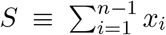 is the number of segregating sites, and *H_n−_*_1_ denotes the *n*-th harmonic number. We then minimize

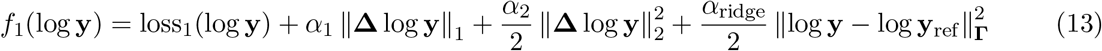

over log **y***∈* ℝ*^m^* to obtain the demographic history. Here, the *α* terms are hyperparameters which we soon describe in more detail.

Having fixed **y** from the previous step, we next infer **Z**. We initialize **Z**= **Z**_ref_ to the MLE constant MuSH: mutation type *j* has the constant rate 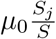, where 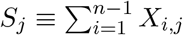 is the number of segregating sites in mutation type *j*. Using the default soft rank penalty, we then minimize

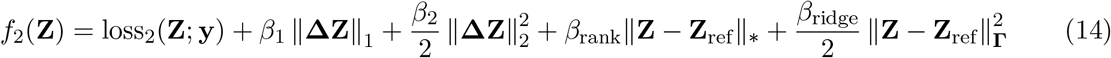

over **Z***∈ R^m×^*^(*κ−*1)^ to obtain the ilr-transformed MuSH. Using the hard rank penalty instead of the default soft rank penalty, we would replace the nuclear norm *||·|| _*_* with the rank function rank(*·*). In equations (13) and (14), the *α* and *β* hyperparameters control the strength of the penalties on **y** and **Z** respectively.

We now briefly cover the methods used for optimization. The cost function (13) is nonconvex due to the nonlinear dependence of ***ξ*** on **y**, while the cost function (14) is convex (although using the hard rank penalty renders it nonconvex). The TV penalties on both (13) and (14) are nonsmooth, as is the soft rank penalty on (14). If the hard rank penalty is used instead of the soft rank penalty, (14) is also nonconvex. Although we cannot guarantee convergence to the global minimum for the demographic history (**y**) problem, we have found that proximal gradient methods rapidly converge to good solutions. Briefly, in proximal methods the cost is split into differentiable and non-differentiable parts, gradient descent steps are taken using the smooth part of the cost, then the *proximal operator* (or *prox*) of the non-differentiable piece is applied. The prox projects to a nearby point which ensures that the nonsmooth part of the cost is small and is easily computed for the TV and hard or soft rank penalties. For the **y** problem, we use the Nesterov accelerated proximal gradient method with adaptive line search [71, 72, 73, 74]. For the MuSH (**Z**) problem, we use a three operator splitting method to deal with the two nonsmooth terms [75]. Our optimization algorithms are implemented very generally as a submodule in the mushi package: https://harrispopgen.github.io/mushi/stubs/mushi.optimization.html. For development purposes, we used similar simulations to those in the main text, but using the mushi forward model instead of msprime (where the PRF likelihood is exact) (see https://harrispopgen.github.io/mushi/developer.html).

### Hyperparameter tuning

Although mushi does not require a parametric model to be specified, it requires the user to tune a few key regularization parameters to target reasonable solutions. This tuning was performed by hand as we now describe. Rather than treat the ridge penalties as adjustable parameters, we fix them by default to *α*_ridge_ = *β*_ridge_ = 10^*−*4^. This leaves the two smoothing parameters *α*_1_ and *α*_2_ for demographic inference. Setting both very small gives erratic, unregularized solutions. Increasing *α*_1_ limits the number of change points, and can be set to produce solutions that are consistent with known features of human demographic history. Subsequently increasing *α*_2_ smooths these change points to produce for example phases of exponential-like growth, but over-smoothing is indicated when the fit to the SFS becomes poor.

We take a similar approach for the MuSH inference step. The three parameters in that case are the smoothing parameters *β*_1_ and *β*_2_ and the complexity parameter *β*_rank_. We set *β*_1_ and *β*_2_ such that most mutation types are nearly flat or smoothly and monotonically varying, while allowing minimal oscillations in mutation types that appear pulse-like in their frequency spectrum (e.g. the TCC*→*TTC pulse). Again, over-smoothing is indicated by poor fit to the *k*-SFS. We set *β*_rank_ to target a specific rank (number of latent histories), generally between 3 and 6. If *β*_rank_ is too large, the rank will be too small to fit all components of the *k*-SFS well. By default we prefer the soft rank penalty for its convexity, but can choose the hard rank penalty if the former results in undesirable shrinkage.

### Software implementation methods

#### The open-source mushi Python package

The mushi software is available as a Python 3 package at https://harrispopgen.github.io/mushi with extensive documentation. We use the JAX package [76] for automatic differentiation and just-in-time compilation of our optimization methods, and the ProxTV package [77] for fast computation of total variation proximal operators. We modified the compositional data analysis module in the scikit-bio package http://scikit-bio.org to allow JAX compatibility. Using default parameters, inferring the demography and MuSH for a population of hundreds of individuals takes a few seconds on a laptop with a modest hardware configuration.

#### Reproducible analysis notebooks

All of the analysis and figures for this paper can be reproduced using Jupyter [78] notebooks available at https://harrispopgen.github.io/mushi. We used msprime [31] and stdpopsim [32] for simulations, TensorLy [79] for NNCP tensor decomposition, umap-learn [44] for UMAP embedding, and several other standard Python packages. We used the Mathematica package fastZeil [81] to procedurally generate recursion formulas for the combinatorial matrix **C** in Theorem 1 (see Appendix A.2).

#### Bioinformatic pipeline for 1000 Genomes Project data

We wrote our pipeline for generating a *k*-SFS for each 1KG population using SCons (https://scons.org), BCFtools (http://samtools.github.io/bcftools), and mutyper (https://github.com/harrispopgen/mutyper). It is available at https://github.com/harrispopgen/mushi/1KG. Pre-computed *k*-SFS data for all 1KG populations is available at https://github.com/harrispopgen/mushi/tree/master/example_data.

1KG variant call data were accessed in BCF format at ftp://ftp.1000genomes.ebi.ac.uk/vol1/ftp/release/20130502/supporting/bcf_files/, with sample manifest available at ftp://ftp.1000genomes.ebi.ac.uk/vol1/ftp/release/20130502/integrated_call_samples_v3.20130502.ALL.panel. Ancestral state estimates were accessed at ftp://ftp.1000genomes.ebi.ac.uk/vol1/ftp/phase1/analysis_results/supporting/ancestral_alignments/human_ancestor_GRCh37_e59, and the strict callability mask was accessed at ftp://ftp.1000genomes.ebi.ac.uk/vol1/ftp/release/20130502/supporting/accessible_genome_masks/20140520.strict_mask.autosomes.bed.

## A Appendix

## A.1 Proof of Theorem 1: the expected SFS given demographic and mutation intensity histories

Suppose *n* haplotypes are sampled in the present, and let random vector **T** = [*T*_2_*,…, T_n_*]^T^ denote the coalescent times measured retrospectively from the present, i.e. *T_n_* is the most recent coalescent time, and *T*_2_ is the TMRCA of the sample.

As in Section 3 of Griffiths and Tavaré [56], we consider a marked Poisson process in which every mutation is assigned a random label drawn iid from the uniform distribution on (0, 1). This is tantamount to the infinite sites assumption, with the unit interval representing the genome, and the random variate labels representing mutant sites. Further suppose that mutation intensity at time *t* (measured retrospectively from the present in units of Wright-Fisher generations) is a function of time 0 *≤ μ*(*t*) *< ∞* (measured in mutations per genome per generation) applying equally to all lines in the coalescent tree. A given line in the coalescent tree then acquires mutations on a genomic subinterval (*p, p* + *dp*) *⊆* (0, 1) at rate *μ*(*t*)*dp*.

Let *ε_dp,b_* denote the event that a mutation present in *b* ∈ {1, 2,…, *n* − 1} haplotypes in the sample occurred within a given genomic interval (*p, p*+*dp*). Given the uniform labeling assumption, the probability of this event is independent of *p*, but the following argument can be generalized to allow the labelling distribution to be nonuniform over the unit interval without changing the result. Let *I_k_* denote the *k*th intercoalescent time interval, i.e. *I_n_* = (0*, T_n_*)*, I_n−_*_1_ = (*T_n_, T_n−_*_1_)*,…, I*_2_ = (*T*_3_*, T*_2_). Let *ε_dp,b,k_* denote the event that the mutation *ε_dp,b_* occurred during interval *I_k_*. For small *dp* and finite *μ*(*t*) we have

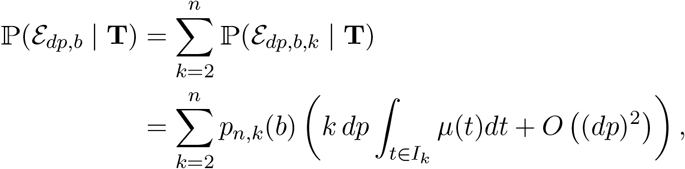

where

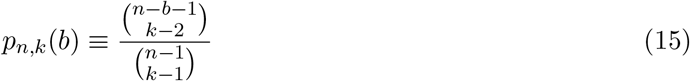

is the probability that a mutant that arose when there were *k* ancestral lines of *n* sampled haplotypes will be present in *b* of them (see [56], eqn. 1.9). The quantity in parentheses is the probability that a mutation arose during the *k*th intercoalescent interval in a genomic interval of size *dp*. Marginalizing **T** gives

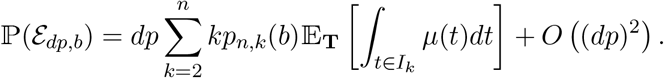

For small *dp*, each genomic interval (*p, p* + *dp*) contains zero or one mutations. Therefore, taking the limit *dp →* 0 and integrating over the genome, the expected number of mutations subtending *b* haplotypes (i.e. the *b*th component of the SFS) is

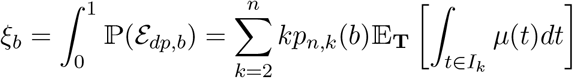

We now substitute in the bounds of every intercoalescent interval *I_k_* = (*T_k_*_+1_*, T_k_*), giving

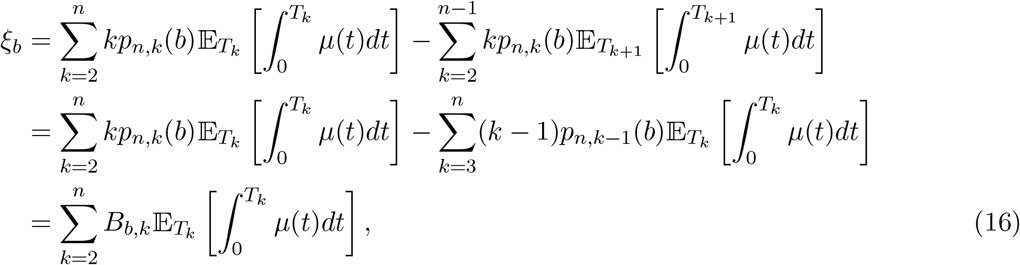

where

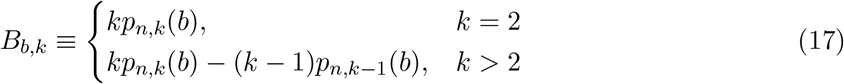

are combinatorial terms.

Polanski et al. [57], eqns. 5-8, give the marginal density for the coalescent time *T_k_* as

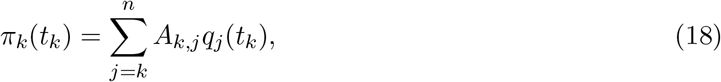

for *k* = 2*,…, n*, where **A** is an (*n −* 1) × (*n −* 1) matrix indexed from 2*,…, n* with

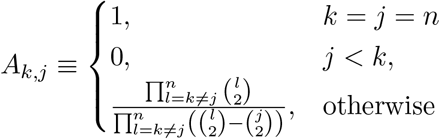

and

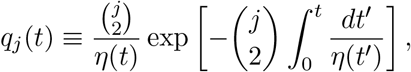

for *j* = 2*,…, n*, and *η*(*t*) is the haploid effective population size history. We assume that 0 *< η*(*t*) *< ∞*. Note that *q_j_*(*t*) is the probability density of the time to the first coalescent event among any subset of *j* individuals in the present, with inhomogeneous Poisson intensity function 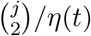.

The expectations in (16) can be expressed using (18) as

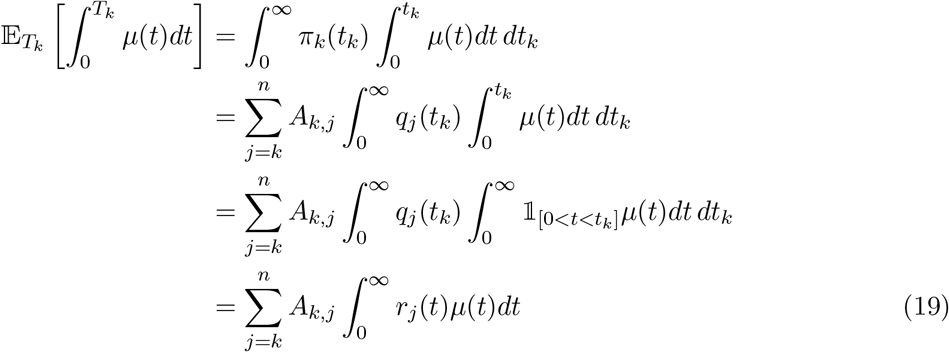

where in the last line we exchange integration order (by Fubini’s theorem) and define the inhomogeneous Poisson survival function

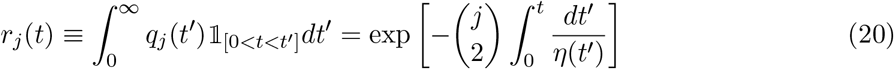

corresponding to density *q_j_*(*t*).

Using (19) in (16) gives

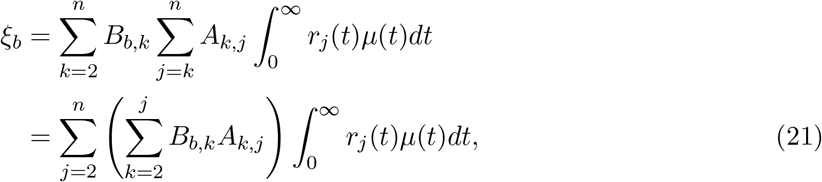

exchanging summation order in the last line. We then have a linear expression for the expected SFS as a function of the mutation intensity history *μ*(*t*):

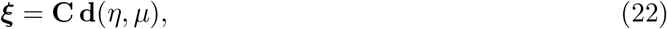

where the (*n −* 1) × (*n −* 1) matrix **C** = **BA** is constant in *μ* and *η*, and

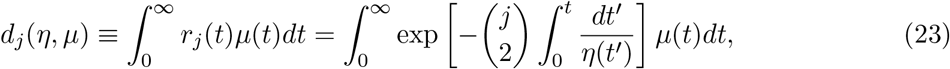

for *j* = 1*,…, n −* 1, is a linear functional of *μ* and a nonlinear functional of *η*.

Given the boundedness assumptions that we have on *η* and *μ*, we now prove boundedness of the map from joint history functions (*η, μ*) to expected SFS vectors *ξ*.

### Lemma 2.

*For all bounded functions η* : ℝ_*≥*0_ *→* ℝ_>0_ *and μ* : ℝ_*≥*0_ *→* ℝ_*≥*0_*, d_j_*(*η, μ*) *is bounded*.

*Proof*. We pass to the diffusion timescale, which measures time in expected number of coalescent events since the present. Let 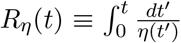, which is strictly increasing ℝ_*≥*0_ *→* ℝ_*≥*0_. Substitute *τ* ≡ *R_η_*(*t*) in (23) to give

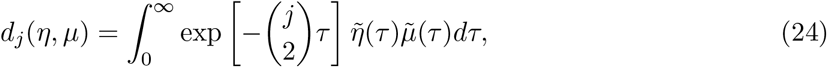

where 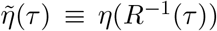 and 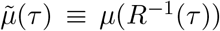. Note that *d_j_* is the Laplace transform of the bounded function 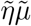 evaluated at 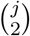, and is thus bounded. In particular,

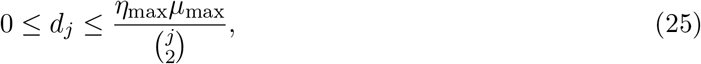

where *η*_max_ and *μ*_max_ are the respective bounds on *η* and *μ*.

The vector **d**(*η, μ*) may be viewed as a nonlinear operator 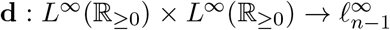 of rank *n −* 1, and is bounded element-wise (Lemma 2). Boundedness of the full operator mapping (*η, μ*) to the expected SFS ***ξ*** follows from the fact that **C** is a matrix with bounded norm. This completes the proof of Theorem 1.

## A.2 Computing the elements of C

We next develop an efficient recursive procedure for computing the matrix **C**. Using (17)

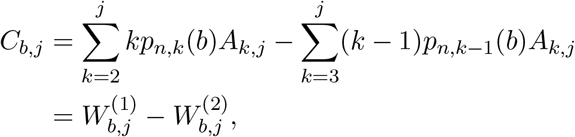

where

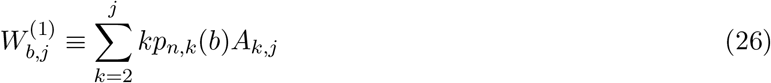

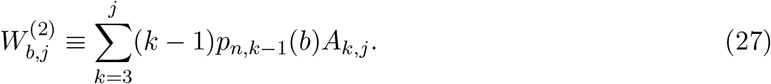

Polanski et al. [58], eqn. 11, show that the nonzero entries of **A** can be expressed as

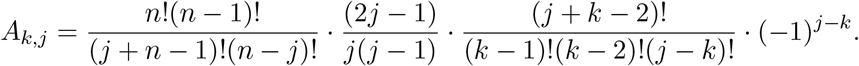

Given the form of *p_n,k_*(*b*) in (15), we see that (26) and (27) are definite sums over hypergeometric terms. We used Zeilberger’s algorithm [59, 81], which finds polynomial recurrences for definite sums of hypergeometric terms, to procedurally generate the following second-order recursions in *j*:

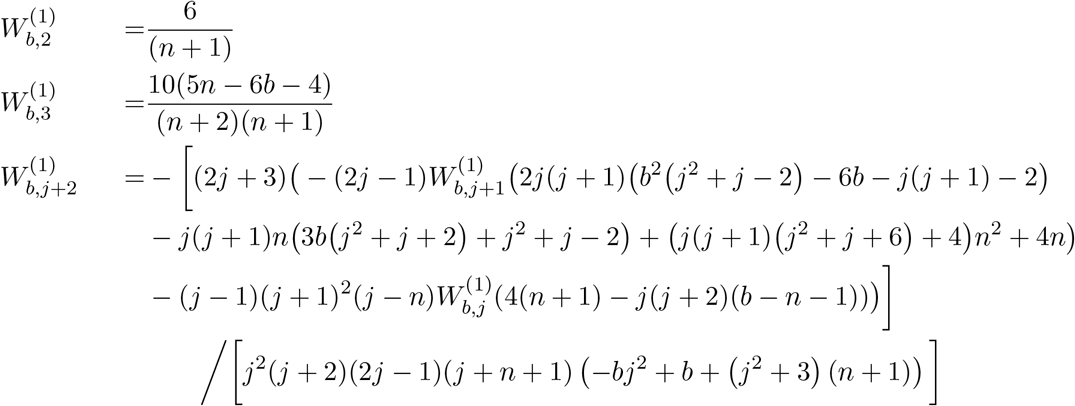

and

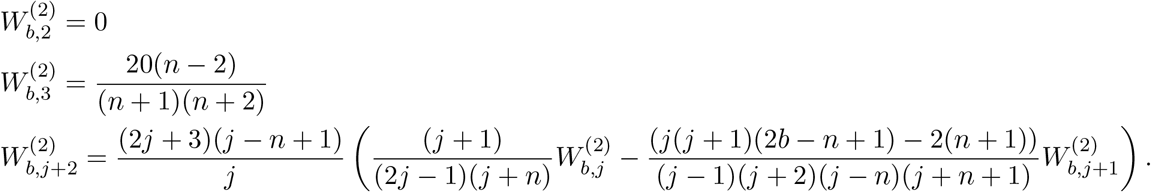

These formulae are used to numerically compute the entries in **C**. The results of this section can be reproduced from the supplementary Mathematica notebook https://github.com/harrispopgen/mushi/blob/master/docsrc/notebooks/recurrence.nb

## A.3 Discretization of history functions and computation of d(*η*, *μ*)

We represent histories as piecewise constant functions of time on *m* pieces [*t*_0_*, t*_1_), [*t*_1_*, t*_2_)*,…,* [*t_m−_*_1_*, t_m_*), where 0 = *t*_0_ *< t*_1_ *< · · · < t_m−_*_1_ *< t_m_* = *∞*. The grid is common to *η*(*t*) and *μ*(*t*). We take the boundaries of the pieces as fixed parameters and in practice use a logarithmically-spaced dense grid of hundreds of pieces to approximate infinite-dimensional histories. Let column vector **y** = [*y*_1_*,…, y_m_*]^T^ denote the constant population size *η*(*t*) during each piece, and let **w** = [*w*_1_*,…, w_m_*]^T^ denote the constant mutation rate *μ*(*t*) during each piece.

With this we can follow the proof of Proposition 1 in [26], *mutatis mutandis*, with our (24) to arrive at

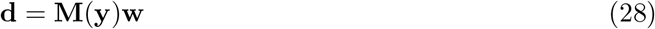

where

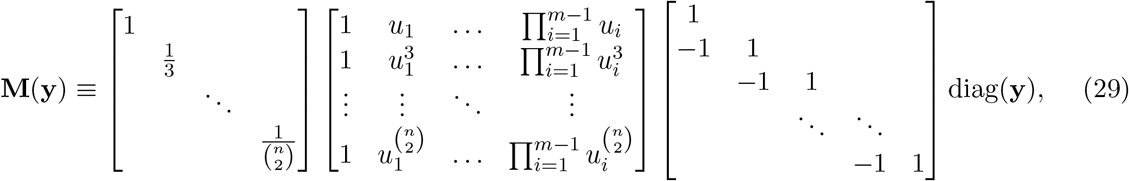

and *u_l_ ≡* exp(*−*(*t_l_ − t_l−_*_1_)*/y_l_*) for *l* = 1*,…, m*. Note that the (*n −* 1) *× m* matrix **M**(**y**) is a nonlinear function of the demographic history **y** because the *u_l_* are nonlinear functions of **y**. This reflects the fact that it is a discretization of the nonlinear operator **d**(*·, μ*). Combining (28) with (22) gives the discretized forward model

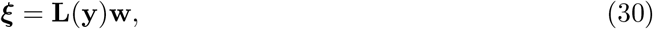

where **L**(**y**) *≡* **CM**(**y**).

## A.4 Proof of Lemma 1

Fix the mutation type *i*, and consider the multinomial over *j*

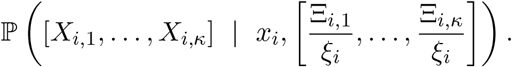

We must show that any element of the multinomial vector

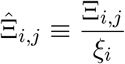

can be formulated without reference to *η*. From elementary properties of the multinomial, the conditional expection value of *X_i,j_* given *x_i_* is

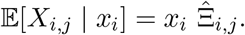

Now, since mutation events are independent we perform a thinning operation on each of the *x_i_* mutation events

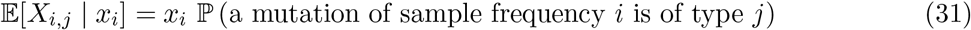

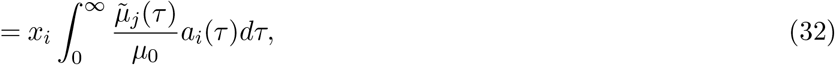

where *a_i_*(*τ*) is the pdf of a mutation’s age *τ* measured in expected coalescent events (diffusion time) conditioned on its sample frequency *i*. So

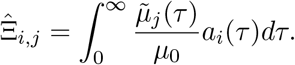

This is independent of *η* by definition of the diffusion time scale as the intensity measure of the coalescent process. This completes the proof of Lemma 1.

## A.5 Tempora incognita: observability toward the coalescent horizon

The time-domain singular vectors of 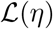 form an eigenbasis for solutions *μ*(*t*) that are possible, in principle, to reconstruct from the SFS. However, sampling noise about the expected SFS will corrupt information from singular vectors that are associated to smaller singular values. These corrupted components will be the directions in solution space associated with higher frequency and less smooth dynamics. Since the singular values of 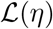 have a very large dynamic range (the condition number is large), the presence of noise will limit reconstruction to smoother, more slowly varying components that are least corrupted and erase information about more sudden events.

Figure 8 depicts the observability of mutation rate history via spectral analysis of 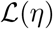 for a case with *η*(*t*) a simple bottleneck history. From (18) and (20) in the Appendix A, the CDF of the TMRCA can be computed given *η*(*t*). We see only the top few components (ranked by singular value) persist at times older than the bottleneck, and all components vanish beyond the TMRCA of the sample. Higher frequency behavior becomes more difficult to observe if it is older than the bottleneck, concretely illustrating how demographic events erase information about population history. The results of this section can be reproduced from the supplementary notebook: https://harrispopgen.github.io/mushi/notebooks/observability.html.

**Figure 8:**
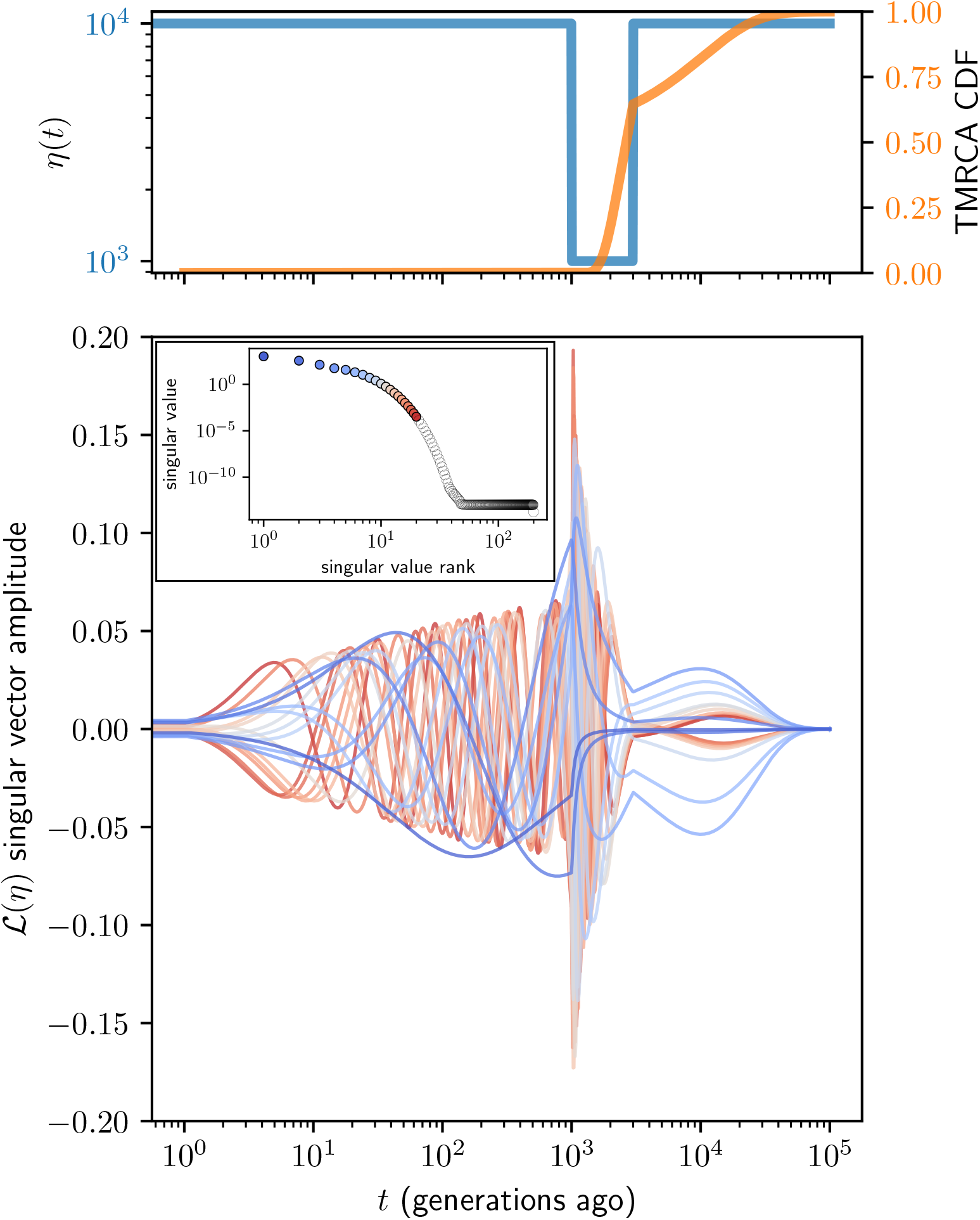
Observability of mutation rate history via the spectral analysis of 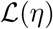 for the case of a bottleneck history. The top panel plots demographic history with a bottleneck from about 3000 to 1000 generations ago (blue, left ordinate), and TMRCA CDF (orange, right ordinate). The bottom panel plots the top 20 time domain singular vectors, with the inset showing the corresponding ranked singular values. Time was discretized with a logarithmic grid of 1000 points, and *n* = 200 sampled haplotypes were assumed.

## References

[1] Kelley Harris. Reading the genome like a history book. Science, 358(6368):1265, December 2017.

[2] J B S Haldane The rate of spontaneous mutation of a human gene. 1935. J. Genet., 83(3): 235–244, December 2004.

[3] Michael W Nachman. Haldane and the first estimates of the human mutation rate. J. Genet., 83(3):231–233, December 2004.

[4] Motoo Kimura. On the evolutionary adjustment of spontaneous mutation rates*. Genet. Res., 9(1):23–34, February 1967.

[5] Michael Lynch, Matthew S Ackerman, Jean-Francois Gout, Hongan Long, Way Sung, W Kelley Thomas, and Patricia L Foster. Genetic drift, selection and the evolution of the mutation rate. Nat. Rev. Genet., 17(11):704–714, October 2016.

[6] M Goodman. Rates of molecular evolution: the hominoid slowdown. Bioessays, 3(1):9–14, July 1985.

[7] Aylwyn Scally and Richard Durbin. Revising the human mutation rate: implications for understanding human evolution. Nat. Rev. Genet., 13(10):745–753, October 2012.

[8] Ziyue Gao, Minyoung J Wyman, Guy Sella, and Molly Przeworski. Interpreting the dependence of mutation rates on age and time. PLoS Biol., 14(1):e1002355, January 2016.

[9] Hákon Jónsson, Patrick Sulem, Birte Kehr, Snaedis Kristmundsdottir, Florian Zink, Eirikur Hjartarson, Marteinn T Hardarson, Kristjan E Hjorleifsson, Hannes P Eggertsson, Sigurjon Axel Gudjonsson, Lucas D Ward, Gudny A Arnadottir, Einar A Helgason, Hannes Helgason, Arnaldur Gylfason, Adalbjorg Jonasdottir, Aslaug Jonasdottir, Thorunn Rafnar, Mike Frigge, Simon N Stacey, Olafur Th Magnusson, Unnur Thorsteinsdottir, Gisli Masson, Augustine Kong, Bjarni V Halldorsson, Agnar Helgason, Daniel F Gudbjartsson, and Kari Stefansson. Parental influence on human germline de novo mutations in 1,548 trios from iceland. Nature, 549(7673):519–522, September 2017.

[10] Raheleh Rahbari, Arthur Wuster, Sarah J Lindsay, Robert J Hardwick, Ludmil B Alexandrov, Saeed Al Turki, Anna Dominiczak, Andrew Morris, David Porteous, Blair Smith, Michael R Stratton, UK10K Consortium, and Matthew E Hurles. Timing, rates and spectra of human germline mutation. Nat. Genet., 48(2):126–133, February 2016.

[11] Dick G Hwang and Phil Green. Bayesian markov chain monte carlo sequence analysis reveals varying neutral substitution patterns in mammalian evolution. Proc. Natl. Acad. Sci. U. S. A., 101(39):13994–14001, September 2004.

[12] Laure Ségurel, Minyoung J Wyman, and Molly Przeworski. Determinants of mutation rate variation in the human germline. Annu. Rev. Genomics Hum. Genet., 15:47–70, June 2014.

[13] Nicolas Galtier and Laurent Duret. Adaptation or biased gene conversion? extending the null hypothesis of molecular evolution. Trends Genet., 23(6):273–277, June 2007.

[14] Laurent Duret and Nicolas Galtier. Biased gene conversion and the evolution of mammalian genomic landscapes. Annu. Rev. Genomics Hum. Genet., 10:285–311, 2009.

[15] Ludmil B Alexandrov, Serena Nik-Zainal, David C Wedge, Samuel A J R Aparicio, Sam Behjati, Andrew V Biankin, Graham R Bignell, Niccol`o Bolli, Ake Borg, Anne-Lise Børresen-Dale, Sandrine Boyault, Birgit Burkhardt, Adam P Butler, Carlos Caldas, Helen R Davies, Christine Desmedt, Roland Eils, Jórunn Erla Eyfjörd, John A Foekens, Mel Greaves, Fumie Hosoda, Barbara Hutter, Tomislav Ilicic, Sandrine Imbeaud, Marcin Imielinski, Natalie Jäger, David T W Jones, David Jones, Stian Knappskog, Marcel Kool, Sunil R Lakhani, Carlos López-Otín, Sancha Martin, Nikhil C Munshi, Hiromi Nakamura, Paul A Northcott, Marina Pajic, Elli Papaemmanuil, Angelo Paradiso, John V Pearson, Xose S Puente, Keiran Raine, Manasa Ramakrishna, Andrea L Richardson, Julia Richter, Philip Rosenstiel, Matthias Schlesner, Ton N Schumacher, Paul N Span, Jon W Teague, Yasushi Totoki, Andrew N J Tutt, Rafael Valdés-Mas, Marit M van Buuren, Laura van ‘t Veer, Anne Vincent-Salomon, Nicola Waddell, Lucy R Yates, Australian Pancreatic Cancer Genome Initiative, ICGC Breast Cancer Consortium, ICGC MMML-Seq Consortium, ICGC PedBrain, Jessica Zucman-Rossi, P Andrew Futreal, Ultan McDermott, Peter Lichter, Matthew Meyerson, Sean M Grimmond, Reiner Siebert, Elías Campo, Tatsuhiro Shibata, Stefan M Pfister, Peter J Campbell, and Michael R Stratton. Signatures of mutational processes in human cancer. Nature, 500(7463):415–421, August 2013.

[16] Thomas Helleday, Saeed Eshtad, and Serena Nik-Zainal. Mechanisms underlying mutational signatures in human cancers. Nat. Rev. Genet., 15(9):585–598, September 2014.

[17] Kelley Harris. Evidence for recent, population-specific evolution of the human mutation rate. Proc. Natl. Acad. Sci. U. S. A., 112(11):3439–3444, March 2015.

[18] Kelley Harris and Jonathan K Pritchard. Rapid evolution of the human mutation spectrum. Elife, 6, April 2017.

[19] Iain Mathieson and David Reich. Differences in the rare variant spectrum among human populations. PLoS Genet., 13(2):e1006581, February 2017.

[20] Jedidiah Carlson, Adam E Locke, Matthew Flickinger, Matthew Zawistowski, Shawn Levy, Richard M Myers, Michael Boehnke, Hyun Min Kang, Laura J Scott, Jun Z Li, Sebastian Zöllner, and BRIDGES Consortium. Extremely rare variants reveal patterns of germline mutation rate heterogeneity in humans. Nat. Commun., 9(1):3753, September 2018.

[21] Ipsita Agarwal and Molly Przeworski. Signatures of replication timing, recombination, and sex in the spectrum of rare variants on the human X chromosome and autosomes. Proc. Natl. Acad. Sci. U. S. A., 116(36):17916–17924, September 2019.

[22] Simon Myers, Charles Fefferman, and Nick Patterson. Can one learn history from the allelic spectrum? Theor. Popul. Biol., 73(3):342–348, May 2008.

[23] Anand Bhaskar and Yun S Song. Descartes’ rule of signs and the identifiability of population demographic models from genomic variation data. Ann. Stat., 42(6):2469–2493, December 2014.

[24] Jonathan Terhorst and Yun S Song. Fundamental limits on the accuracy of demographic inference based on the sample frequency spectrum. Proc. Natl. Acad. Sci. U. S. A., 112(25):7677–7682, June 2015.

[25] Soheil Baharian and Simon Gravel. On the decidability of population size histories from finite allele frequency spectra. Theor. Popul. Biol., 120:42–51, March 2018.

[26] Zvi Rosen, Anand Bhaskar, Sebastien Roch, and Yun S Song. Geometry of the sample frequency spectrum and the perils of demographic inference. Genetics, page genetics.300733.2018, July 2018.

[27] 1000 Genomes Project Consortium, Adam Auton, Lisa D Brooks, Richard M Durbin, Erik P Garrison, Hyun Min Kang, Jan O Korbel, Jonathan L Marchini, Shane McCarthy, Gil A McVean, and Gonçalo R Abecasis. A global reference for human genetic variation. Nature, 526(7571):68–74, October 2015.

[28] Leo Speidel, Marie Forest, Sinan Shi, and Simon R Myers. A method for genome-wide genealogy estimation for thousands of samples. Nat. Genet., 51(9):1321–1329, September 2019.

[29] J K Pritchard, M Stephens, and P Donnelly. Inference of population structure using multilocus genotype data. Genetics, 155(2):945–959, June 2000.

[30] S A Sawyer and D L Hartl. Population genetics of polymorphism and divergence. Genetics, 132(4):1161–1176, December 1992.

[31] Jerome Kelleher, Alison M Etheridge, and Gilean McVean. Efficient coalescent simulation and genealogical analysis for large sample sizes. PLoS Comput Biol, 12(5):1–22, 05 2016. doi: 10.1371/journal.pcbi.1004842. URL http://dx.doi.org/10.1371%2Fjournal.pcbi.1004842.

[32] Jeffrey R Adrion, Christopher B Cole, Noah Dukler, Jared G Galloway, Ariella L Gladstein, Graham Gower, Christopher C Kyriazis, Aaron P Ragsdale, Georgia Tsambos, Franz Baumdicker, Jedidiah Carlson, Reed A Cartwright, Arun Durvasula, Bernard Y Kim, Patrick McKenzie, Philipp W Messer, Ekaterina Noskova, Diego Ortega-Del Vecchyo, Fernando Racimo, Travis J Struck, Simon Gravel, Ryan N Gutenkunst, Kirk E Lohmeuller, Peter L Ralph, Daniel R Schrider, Adam Siepel, Jerome Kelleher, and Andrew D Kern. A community-maintained standard library of population genetic models. December 2019.

[33] International HapMap Consortium. A second generation human haplotype map of over 3.1 million SNPs. Nature, 449(7164):851–861, October 2007.

[34] Stephan Schiffels and Richard Durbin. Inferring human population size and separation history from multiple genome sequences, 2014.

[35] Jonathan Terhorst, John A Kamm, and Yun S Song. Robust and scalable inference of population history from hundreds of unphased whole genomes. Nat. Genet., 49(2):303–309, February 2017.

[36] Jonathan G Terhorst. Demographic Inference from Large Samples: Theory and Methods. PhD thesis, UC Berkeley, 2017.

[37] Heng Li and Richard Durbin. Inference of human population history from individual whole-genome sequences. Nature, 475(7357):493–496, July 2011.

[38] Jeffrey P Spence, John A Kamm, and Yun S Song. The site frequency spectrum for general coalescents. Genetics, 202(4):1549–1561, April 2016.

[39] Aylwyn Scally. The mutation rate in human evolution and demographic inference. Curr. Opin. Genet. Dev., 41:36–43, December 2016.

[40] Jack N Fenner. Cross-cultural estimation of the human generation interval for use in genetics-based population divergence studies, 2005.

[41] Jacob A Tennessen, Abigail W Bigham, Timothy D O’Connor, Wenqing Fu, Eimear E Kenny, Simon Gravel, Sean McGee, Ron Do, Xiaoming Liu, Goo Jun, Hyun Min Kang, Daniel Jordan, Suzanne M Leal, Stacey Gabriel, Mark J Rieder, Goncalo Abecasis, David Altshuler, Deborah A Nickerson, Eric Boerwinkle, Shamil Sunyaev, Carlos D Bustamante, Michael J Bamshad, Joshua M Akey, Broad GO, Seattle GO, and NHLBI Exome Sequencing Project. Evolution and functional impact of rare coding variation from deep sequencing of human exomes. Science, 337(6090):64–69, July 2012.

[42] Tamara G Kolda and Brett W Bader. Tensor decompositions and applications. SIAM Rev., 51(3):455–500, August 2009.

[43] Luke Anderson-Trocmé, Rick Farouni, Mathieu Bourgey, Yoichiro Kamatani, Koichiro Higasa, Jeong-Sun Seo, Changhoon Kim, Fumihiko Matsuda, and Simon Gravel. Legacy data confounds genomics studies. Mol. Biol. Evol., August 2019.

[44] Leland McInnes, John Healy, and James Melville. UMAP: Uniform manifold approximation and projection for dimension reduction. February 2018.

[45] Michael E Goldberg and Kelley Harris. Great ape mutation spectra vary across the phylogeny and the genome due to distinct mutational processes that evolve at different rates. October 2019.

[46] Beth L Dumont. Significant strain variation in the mutation spectra of inbred laboratory mice. Mol. Biol. Evol., 36(5):865–874, May 2019.

[47] Berit Lindum Waltoft and Asger Hobolth. Non-parametric estimation of population size changes from the site frequency spectrum. Stat. Appl. Genet. Mol. Biol., 17(3), June 2018.

[48] Ryan J Tibshirani. Adaptive piecewise polynomial estimation via trend filtering. Ann. Stat., 42(1):285–323, February 2014.

[49] Annabel C Beichman, Tanya N Phung, and Kirk E Lohmueller. Comparison of single genome and allele frequency data reveals discordant demographic histories. G3, 7(11):3605–3620, November 2017.

## References

[50] J F C Kingman. The coalescent. Stochastic Process. Appl., 13(3):235–248, September 1982.

[51] J F C Kingman. On the genealogy of large populations. J. Appl. Probab., 19(A):27–43, 1982.

[52] J F C Kingman. Exchangeability and the evolution of large populations, exchangeability in probability and statistics (rome, 1981), 1982.

[53] J F Kingman. Origins of the coalescent. 1974-1982. Genetics, 156(4):1461–1463, December 2000.

[54] John Wakeley. Coalescent theory: an introduction. 2009.

[55] Warren J Ewens. Mathematical Population Genetics 1: Theoretical Introduction. Springer Science & Business Media, October 2012.

[56] R C Griffiths and S Tavaré. The age of a mutation in a general coalescent tree. Stoch. Models, 1998.

[57] A Polanski, A Bobrowski, and M Kimmel. A note on distributions of times to coalescence, under time-dependent population size. Theor. Popul. Biol., 63(1):33–40, February 2003.

[58] A Polanski and M Kimmel. New explicit expressions for relative frequencies of single-nucleotide polymorphisms with application to statistical inference on population growth. Genetics, 165 (1):427–436, September 2003.

[59] Marko Petkovšek, Herbert S Wilf, and Doron Zeilberger. A= b, ak peters ltd. Wellesley, MA, 30, 1996.

[60] Jason Schweinsberg. Coalescents with simultaneous multiple collisions. Electron. J. Probab., 5, 2000.

[62] J Aitchison. The statistical analysis of compositional data. J. R. Stat. Soc. Series B Stat. Methodol., 44(2):139–160, January 1982.

[63] Vera Pawlowsky-Glahn, Juan José Egozcue, and Raimon Tolosana-Delgado. Modeling and Analysis of Compositional Data. John Wiley & Sons, March 2015.

[64] Charles L Epstein and John Schotland. The bad truth about laplace’s transform. SIAM Rev., 50(3):504–520, January 2008.

[65] Trevor Hastie, Robert Tibshirani, and Martin Wainwright. Statistical Learning with Sparsity: The Lasso and Generalizations. CRC Press, May 2015.

[66] Robert Tibshirani, Michael Saunders, Saharon Rosset, Ji Zhu, and Keith Knight. Sparsity and smoothness via the fused lasso. J. R. Stat. Soc. Series B Stat. Methodol., 67(1):91–108, February 2005.

[67] Leonid I Rudin, Stanley Osher, and Emad Fatemi. Nonlinear total variation based noise removal algorithms. Physica D, 60(1–4):259–268, 1992.

[68] Grace Wahba. Spline Models for Observational Data. SIAM, January 1990.

[69] M Fazel, H Hindi, and S P Boyd. A rank minimization heuristic with application to minimum order system approximation. In Proceedings of the 2001 American Control Conference. (Cat. No.01CH37148), volume 6, pages 4734–4739 vol.6, June 2001.

[70] Anirban DasGupta. Probability for statistics and machine learning : fundamentals and advanced topics. Springer texts in statistics. Springer, New York, 2011. ISBN 9781441996343.

[71] YE Nesterov. A method for solving the convex programming problem with convergence rate o(1*/k*^2^). Dokl. Akad. Nauk SSSR, 269:543–547, 1983.

[72] Yurii Nesterov. Lectures on Convex Optimization. Springer International Publishing, December 2018.

[73] Paul Tseng. On accelerated proximal gradient methods for convex-concave optimization. submitted to SIAM Journal on Optimization, 2:3, 2008.

[74] Amir Beck and Marc Teboulle. A fast iterative Shrinkage-Thresholding algorithm for linear inverse problems. SIAM J. Imaging Sci., 2(1):183–202, January 2009.

[75] Fabian Pedregosa and Gauthier Gidel. Adaptive three operator splitting. April 2018.

[76] James Bradbury, Roy Frostig, Peter Hawkins, Matthew James Johnson, Chris Leary, Dougal Maclaurin, and Skye Wanderman-Milne. JAX: composable transformations of Python+NumPy programs, 2018. URL http://github.com/google/jax.

[77] Álvaro Barbero and Suvrit Sra. Modular proximal optimization for multidimensional total-variation regularization. November 2014.

[78] Thomas Kluyver, Benjamin Ragan-Kelley, Fernando Pérez, Brian E Granger, Matthias Bussonnier, Jonathan Frederic, Kyle Kelley, Jessica B Hamrick, Jason Grout, Sylvain Corlay, Paul Ivanov, Damián Avila, Safia Abdalla, Carol Willing, and others. Jupyter notebooks - a publishing format for reproducible computational workflows. ELPUB, 2016.

[79] Jean Kossaifi, Yannis Panagakis, Anima Anandkumar, and Maja Pantic. TensorLy: Tensor learning in python. October 2016.

[81] Peter Paule and Markus Schorn. A mathematica version of zeilberger’s algorithm for proving binomial coefficient identities. Journal of symbolic computation, 20(5-6):673–698, 1995.

